# Gene amplifications cause high-level resistance against albicidin in Gram-negative bacteria

**DOI:** 10.1101/2022.09.15.507240

**Authors:** Mareike Saathoff, Simone Kosol, Torsten Semmler, Karsten Tedin, Nicole Dimos, Johannes Kupke, Maria Seidel, Fereshteh Ghazisaeedi, Silver A. Wolf, Benno Kuropka, Wojciech Czyszczoń, Dmitry Ghilarov, Stefan Grätz, Jonathan G. Heddle, Bernhard Loll, Roderich D. Süssmuth, Marcus Fulde

## Abstract

Antibiotic resistance is a continuously increasing concern for public health care. Understanding resistance mechanisms and their emergence is crucial for the development of new antibiotics and their effective use. Here, we report the discovery of a gene amplification-based mechanism that imparts an up to 1000-fold increase in resistance levels against the antibiotic albicidin. We show that this mechanism protects *Salmonella* Typhimurium and *Escherichia coli* by increasing the copy number of the GyrI-like transcription regulator STM3175 (YgiV) which binds albicidin. X-ray crystallography and molecular docking studies reveal a conserved binding motif that can interact with aromatic building blocks of albicidin. Phylogenetic studies suggest that this resistance mechanism is ubiquitous in Gram-negative bacteria and our experiments confirm that STM3175 homologs can convey resistance in pathogens such as *Vibrio vulnificus* and *Pseudomonas aeruginosa*.

## Introduction

Antimicrobial resistance (AMR) in pathogenic, commensal and food-borne bacteria remains a major health hazard for humans. According to a recent study, in 2019 almost 5 million human deaths were estimated to be associated with bacterial AMR, with a prognosis of an increase of up to 10 million deaths by 2050. It is widely accepted, that inappropriate use of antimicrobials as therapeutics and feed additives in human and veterinary medicine coupled with a lack in understanding of bacterial resistance mechanisms account for the worldwide increase of resistant and multi-resistant bacteria (*1, 2*). To develop effective long-term solutions to combat antimicrobial resistance and improve public health, understanding resistance mechanisms and their evolution is crucial.

Bacterial adaptation to stress conditions occurs via different types of responses. In addition to stable genetic, *bona fide* resistance mechanisms, transient and unstable responses conferred, e.g. by differential gene regulation and protein stability, and changes in gene copy numbers, so-called gene duplication-amplification (GDA) events, are frequently observed (*3–5*). The latter occur rapidly (e.g. <10 generations), whereas establishing genetic modifications takes thousands of generations. GDAs have been observed as a consequence of selective pressure from antibiotic exposure, often resulting in elevated cellular concentrations of a modifying or degrading enzyme, or of efflux pumps. In addition to their role in adapting bacteria quickly to harmful conditions, GDAs have also been implicated in the evolution of *bona fide* resistance mechanisms on a populational (6) and on a sub-populational level (so called heteroresistance) (7). However, as knowledge about the occurrence of GDAs in antibiotic treatment is still scarce and their verification in routine diagnostics is difficult, it must be assumed that GDAs have a non-negligible impact on the success of antibiotic treatment of infectious diseases.

The promising antibacterial peptide albicidin was first isolated as a phytotoxic compound from the plant-pathogenic bacterium *Xanthomonas albilineans* (*6, 7*). Since then, albicidin (*8–10*) as well as the related compounds cystobactamids (*11, 12*) and coralmycins (*13, 14*) are undergoing development towards clinical use. Albicidin is active at nanomolar concentrations against a range of different Gram-positive and Gram-negative bacterial species, including *Klebsiella aerogenes*, *Salmonella* Typhimurium, *Escherichia coli*, as well as the ESKAPE pathogens *Enterococcus faecium, Staphylococcus aureus*, *Klebsiella pneumonia*, *Acinetobacter baumanii*, *Pseudomonas aeruginosa*, and *Enterbacter spp.*. The antibiotic peptide inhibits the supercoiling activity of bacterial DNA gyrase (topoisomerase II) and effectively traps the covalent complex formed between DNA and gyrase at concentrations below those of most coumarin antibiotics and quinolones (*15, 16*). Several albicidin resistance mechanisms have been described, e.g. the ABC transporter AlbF that conveys auto-resistance in *Xanthomonas* via the active efflux of albicidin (*17*). Further strategies comprise albicidin degradation by the endopeptidase AlbD produced by *Pantoea dispersa* by specifically cleaving a benzamide bond (*18*) or albicidin binding through the MerR-like transcriptional regulator AlbA synthesized by *Klebsiella oxytoca* (*19*). In *E*. *coli*, adaption to albicidin is frequently linked to mutations in the nucleoside-specific transporter Tsx, resulting in the loss of active transport of albicidin across the outer membrane (*20, 21*).

Here, we report that exposure of *Salmonella* Typhimurium and *Escherichia coli* to increasing concentrations of the gyrase poison albicidin results in chromosomal duplication-amplification. The affected region harbors the GyrI-like domain containing transcription regulator STM3175 (YgiV in *E. coli*), which we identify as a critical factor for conferring high-level resistance against albicidin. We further illustrate that this resistance mechanism is ubiquitous in Gram-negative bacterial species with STM3175 homologs conferring resistance in *Vibrio* and *Pseudomonas*.

## Results

### Exposure to increasing albicidin concentration leads to Tsx transporter mutations and results in gene duplications

To investigate the evolution of albicidin resistance in *S*. Typhimurium, we exposed the wild type (WT) strain to albicidin concentrations considerably higher than the MIC (0.06 μg ml^−1^) and monitored growth over a 24 h period. Because of its higher stability, we used the azahistidine analog of albicidin (Fig. 1A) which has similar activity compared to natural albicidin (*8*). 14 out of 90 clones tested showed growth under these conditions (Fig. S1). Genome sequencing of these albicidin resistant strains revealed that the majority (9 strains) carried mutations in the *tsx* gene. These mutations were associated with a 16-fold increased MIC, compared to the WT, similar to the isogenic *tsx*-deficient control strain (Figure 1A, Table S1). Remarkably, one of the remaining five resistant strains, with an intact *tsx* gene (strain S41), exhibited a more than 100-fold elevated MIC and was able to tolerate concentrations as high as 2 μg mL^−1^ albicidin (Figure 1A). Analysis of DNA sequencing data of strain S41 revealed a gene amplification resulting in three to four copies of a ~47 kb genomic region but no single nucleotide polymorphisms (SNP) (Table S1, S2).

**Fig. 1.**
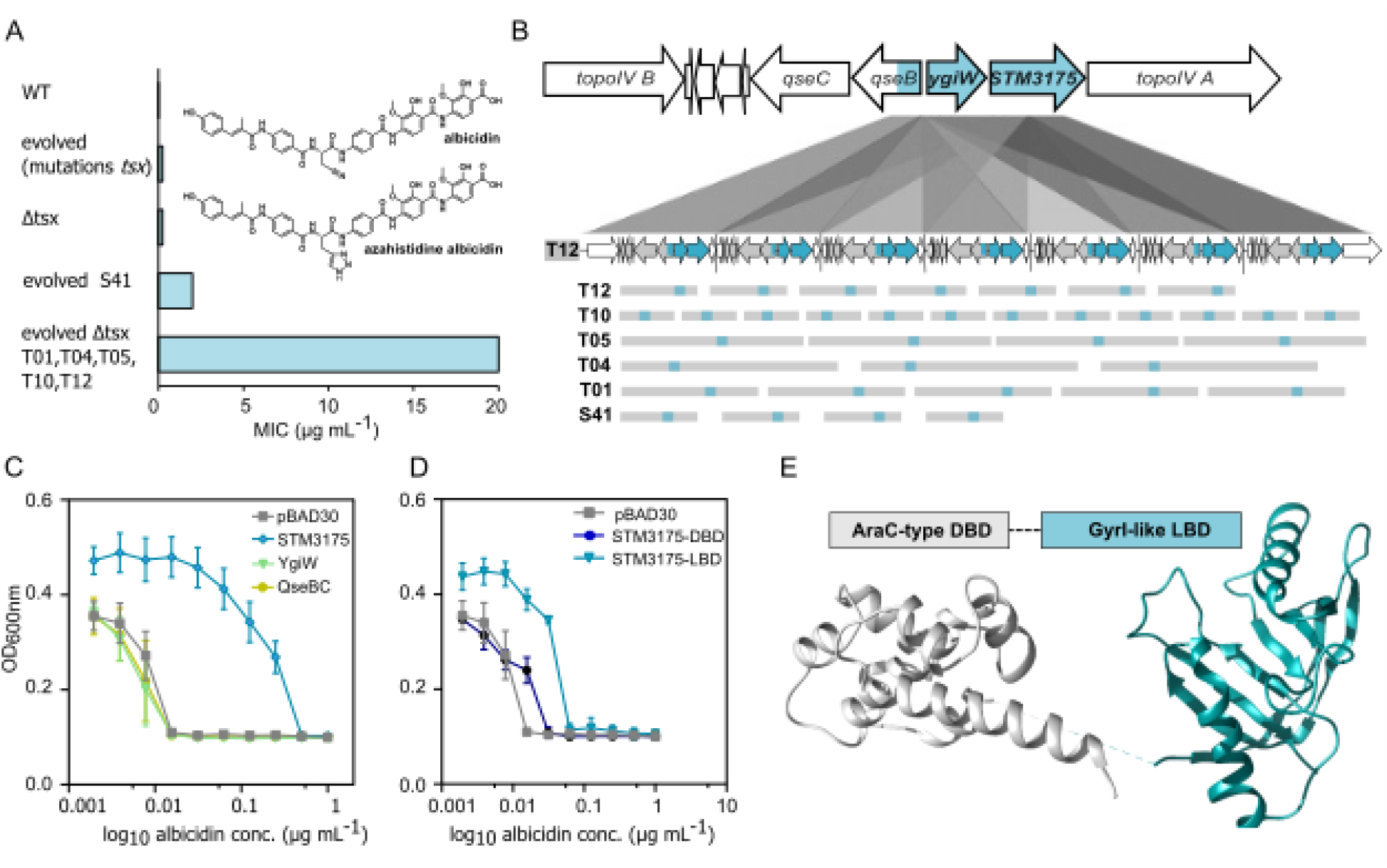
Albicidin high-level resistance resulting from GDAs in evolved *S*. Typhimurium. A) Comparison of albicidin MICs for WT and evolved *S*. Typhimurium strains (see Figure S1, Table S2). A structure of albicidin and azahistidine albicidin is included in the bar diagram. B) Nanopore sequencing revealed 7 copies of the GDA region in the evolved *tsx*-deficient *S*. Typhimurium strain T12, (see below, T12) including the genes between topoisomerase IV subunits A and B (see above, WT). The common ~2200 bp long region present in all six evolved strains (S41, T01, T04, T05, T10, T12) is highlighted in cyan. It includes three genes: *STM3175, ygiW* and partly *qseB*. C) Arabinose-induced over-expression of STM3175 (cyan), but not YgiW (light green) or QseBC olive), results in an elevated albicidin MIC in *S*. Typhimurium WT. D) Arabinose-induced overexpression of the ligand-binding (LBD, cyan), but not of the DNA-binding domain (DBD, blue) of STM3175 revealed and increased MIC against albicidin in *S*. Typhimurium. E) Crystal structure of STM3175 with the conserved AraC-type DBD and GyrI-like LBD shown in grey and cyan, respectively (please also see Fig. S11).

The GDAs caught our interest and we decided to investigate this mechanism in more detail. We therefore repeated the evolution experiments with a mutant strain lacking the *tsx* gene (Δ*tsx*) in order to avoid resistance effects resulting from mutations in the nucleoside transporter gene. After exposing the Δ*tsx* strain stepwise to increasing albicidin concentrations from 0.125 μg ml^−1^ to 20 μg ml^−1^ (Fig. 1A), similar genome alterations were observed as in the *tsx-*proficient WT strains that adapted to the antibiotic. Whole genome sequencing revealed that five out of ten tested strains harbored GDAs ranging between 3 kb to 158 kb (strains T01, T04, T05, T10, T12; Fig. S2A, Table S2) with copy numbers varying between 3 and 15. In these strains, either no SNPs (2/5: T01, T10) or SNPs in genes concerning the common segment of the GDA-region (3/5: T04, T05, T12 - *qseB, stm3175* or *topoisomerase IV subunit B* and additionally one hypothetical protein (1/5: T12)) were identified (Table S3), suggesting that this particular GDA-region is responsible for the significantly increased albicidin tolerance in *S*. Typhimurium: 80-fold compared to the input strain or more than 1000-fold compared to the wild-type strain (Fig. 1A). As mentioned above, a hallmark of GDAs is their reversibility in the absence of the appropriate selection pressure. To test, we randomly selected one of the evolved strains (T12) and incubated it as biological triplicates for 24 hours over a period of 15 days (equivalent to approximately 1500 generations) without albicidin. As shown in Table S4, we demonstrated a reduction in gene duplications in one of the three cases, which correspondingly resulted in a lower MIC to albicidin. The other clones showed no changes in the number of GDAs. However, it is widely known that the phenomenon of heteroresistance is a complex mechanism that happens at the single cell level and in which phenomena such as compensation and biologic fitness play a role, further experiments are needed to elucidate in detail the genesis of albicidin-mediated GDAs.

### GDAs contain genes for the sensory histidine kinase system QseBC and the regulatory protein STM3175

Alignments of the GDAs revealed a conserved ~2200 bp overlap between the amplified regions present in all six (S41, T01, T04, T05, T10, T12) analyzed strains (Fig. S2). The overlapping region contains the two genes *STM3175* and *ygiW*, that are transcribed polycistronically in an operon structure as well as the N-terminal part of *qseB* (Fig. 1B, S2B). Interestingly, *STM3175* and *ygiW* are located directly upstream of the gene encoding DNA topoisomerase IV subunit A (Fig. 1B, S2B). Not much is known about the function of YgiW, a putative periplasmic protein, or STM3175, a putative regulatory protein. QseB and the adjacently encoded QseC (Fig. 1B) belong to a known two-component quorum sensing system consisting of a sensory histidine kinase (QseC) and its response regulator (QseB). To investigate if the absence of these genes affects albicidin tolerance of *S*. Typhimurium, we constructed knockout mutants of *qseBC* and *STM3175-ygiW* of the WT and Δ*tsx* strain. In MIC assays, no detectable significant change in albicidin sensitivity was observed compared to the parent strains (Fig. S3), suggesting that these genes only affect albicidin tolerance in GDAs and higher copy numbers.

### GDAs of the regulator STM3175 afford albicidin resistance

To investigate which of the amplified genes is responsible for the increased albicidin tolerance, MIC assays with *STM3175, ygiW* and *qseB* were conducted under control of an arabinose-inducible promoter. Each gene was cloned into the low-copy number plasmid pBAD30 and expressed in *S*. Typhimurium WT cells. After induction with arabinose, only STM3175 imparted resistance, whereas over-expression of YgiW and QseB did not elicit albicidin tolerance (Fig. 1C and S4 A-B). Notably, the level of albicidin resistance increased with increasing arabinose concentration (Fig. S5). This suggested that the resistance mechanism solely depended on the increase of STM3175 concentration, which would be elevated by the increased copy number of STM3175 in the GDAs. Indeed, RNA-Seq analysis showed a clear increase in mRNA levels of the *STM3175-ygiW* operon as well as the other constituents of the GDA, including *qseB/C* but not the flanking topoisomerase IV subunits A and B (Fig. S6). Proteomic analyses further confirmed higher levels of STM3175, YgiW and QseB/C in the mutant T12 compared to WT cells (Fig. S6, Table S13).

Closer inspection of its amino acid sequence revealed that STM3175 consists of two domains: an N-terminal AraC-like DNA-binding domain (DBD), and a C-terminal GyrI-like ligand-binding domain (LBD). In over-expression experiments, the LBD but not the DBD of STM3175 conveyed resistance (Fig. 1D and S4 C-D). Interestingly, the full-length protein appeared to confer higher resistance than the LBD alone. Further even expression of the LBD in a STM3175 mutant strain, showed the same resistance level as expressed in WT cells with intact STM3175 (Table S5). In agar diffusion assays, both the full-length protein and the LBD alone neutralized the effects of albicidin (Fig. S7). When a two-fold excess of albicidin was added to the STM3175 or LBD proteins before spotting, bacterial growth was clearly reduced, albeit less severely than in the control sample without protein (Fig. S7 C and E), excluding enzymatic modification or degradation of albicidin as mechanism of action for STM3175.

### STM3175 structure

The dual domain structure of a helix-turn-helix DNA-binding element combined with an effector binding domain is reminiscent of AlbA (*19*) (Fig. S8A), the MerR-like transcription factor that conveys albicidin resistance in *Klebsiella oxytoca*. However, the affinity for albicidin and the dual domain structure is where the similarities end. Not only is the sequence identity low (15 %; Fig. S8B) but the predicted secondary structure of the ligand-binding domains is markedly different. AlbA exclusively consists of helical elements whereas STM3175 is composed of α-helices and β-sheets (Fig. S9). CD spectroscopy of recombinantly expressed proteins confirmed the mixed content of α-helix and β-sheet in STM3175 with clearly reduced helical contributions when the DNA-binding domain is not present (Fig. S10 A-C).

This is in excellent agreement with our crystal structure of full-length STM3175 (PDB-ID: 7R3W) where the N-terminal domain consists of seven helices that form two helix-turn-helix DNA-binding motifs typically found in AraC DNA-binding domains (*23*) (Fig. 1E and Fig. S11). The DNA-binding domain is connected to the C-terminal LBD domain via a short helical element and a short linker (~10 aa) which was not resolved in the electron density. The ligand binding domain shows the characteristic GyrI-like fold of two SH2 motifs with a pseudo two-fold symmetry forming a central groove which is clearly visible in the crystal structure despite the relatively low resolution of 3.6 Å (Fig. 1E). The dimensions of the groove (length ~ 30 Å, width ~ 10 Å, height ~ 12 Å) provide an ideal environment for accommodating compounds of linear architecture (*24*). Despite presenting as a monomer in analytical gel filtration (Fig. S12), STM3175 forms domain-swapped dimers in the crystal, where the DBD of one polypeptide forms contacts with the LBD of a second molecule (Fig.S13A-C). The relative orientation of the N-and C-terminal domains of the two molecules is not identical, resulting in an asymmetrical dimer (Fig. S12D, E).

When generating homology models by RoseTTAFold (*25*) and Phyre2 (*26*), the *E. coli* transcription factor Rob (PDB-ID: 1d5y) (*27*) and GyrI (PDB-ID: 1jyha) (*24*) were identified as highest-ranking structural homologs (100 and 99.9% confidence, respectively; Fig. S14). While the structures of the N- and C-terminal domains in the crystal structure and homology model were very similar (RMSD < 1.5 Å), the relative domain orientation differed significantly. The homology models showed a more compact structure, similar to that of Rob bound to DNA (Fig. S14A). This flexibility in the relative domain orientation is consistent with STM3175 structure models by AlphaFold (*28, 29*) (model available in AlphaFold Protein Structure Database under UniProt ID: Q8ZM00), where the predicted aligned error plot also suggests lower confidence in inter-domain accuracy of the prediction. Rob and GyrI, as well as a number of other GyrI-like domain containing proteins, are also found when submitting the STM3175 crystal structure to the DALI server (*30*) to identify structurally similar proteins in the PDB. However, in contrast to other Gyrl-homologs, STM3175 lacks the highly conserved Glu residue in the center of the binding groove, and the charges in and around the groove are not exclusively negative as in most GyrI-like domains (*31*). In STM3175, the groove is mostly lined with hydrophobic residues and extends into a tunnel towards the C-terminus of the LBD (Fig. S11).

### STM3175 albicidin binding

Since several Trp residues are located in and around the binding groove, we monitored the interaction with albicidin by tryptophan fluorescence quenching experiments. To obtain affinity constants, albicidin was titrated to STM3175 or STM3175-LBD and the concentration dependent decrease in the fluorescence signal was measured. Fitting of the quenching data suggests that STM3175 binds albicidin with sub-micromolar affinity (Kd = 0.17 ± 0.01 μM, Fig. 2A). The ligand-binding domain alone binds with similar but somewhat lower affinity (Kd = 0.35 ± 0.05 μM, Fig. 2B) suggesting that the DNA-binding domain is not or only little involved in albicidin binding.

**Fig. 2.**
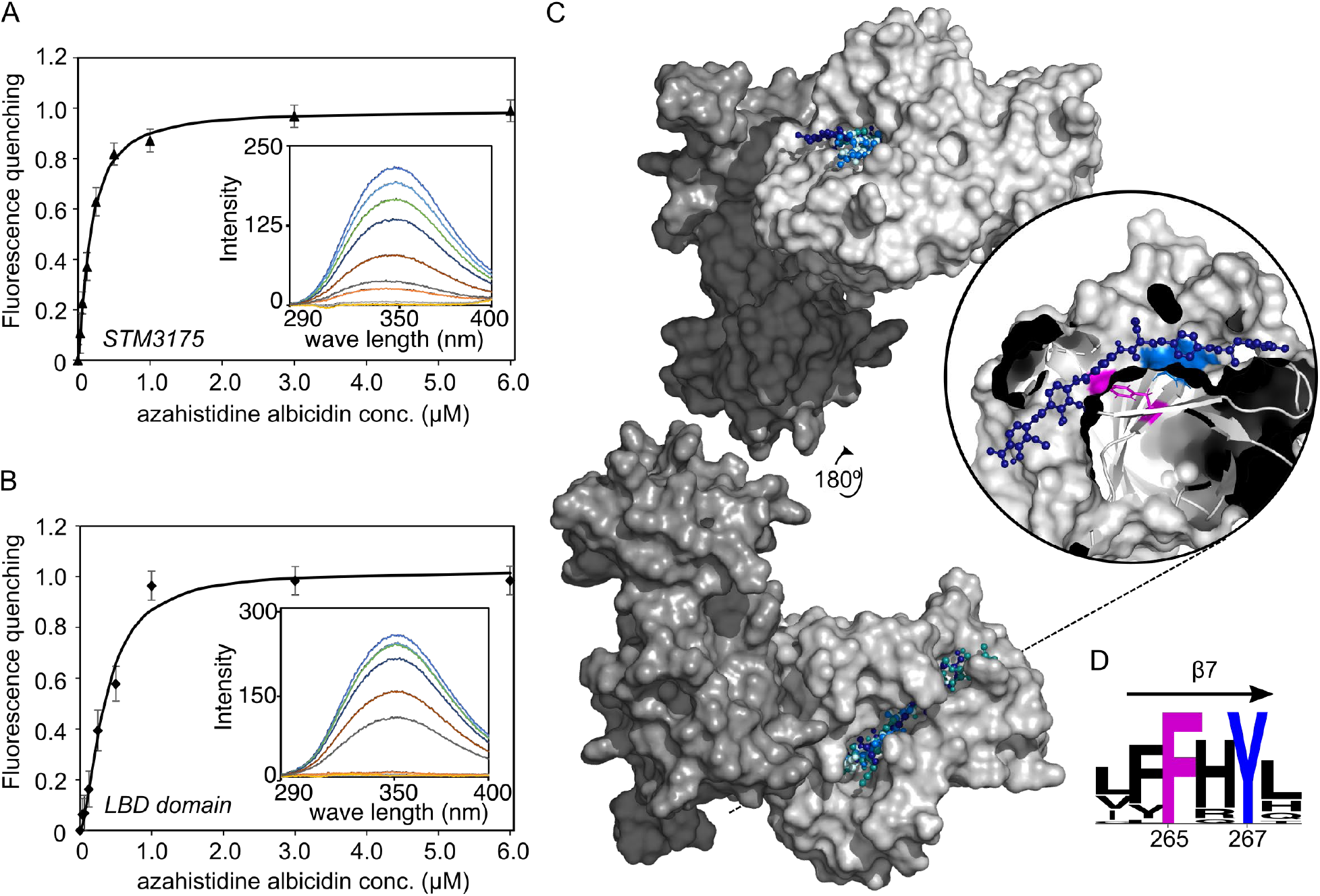
Albicidin binding to STM3175. The binding affinity of STM3175 and its ligand-binding domain (LBD) were determined by monitoring quenching of tryptophan fluorescence emission of STM3175 after addition of azahistidine albicidin. A) Fluorescence quenching of STM3175 plotted as a function of increasing azahistidine albicidin concentration. Fitting of the decreasing fluorescence emission yielded a Kd of 0.17 ± 0.01 μM. B) Fluorescence quenching of STM3175-LBD plotted against increasing azahistidine albicidin concentrations. Fitting of the data yielded a Kd of 0.35 ± 0.05 μM. Insets show emission spectra of STM3175 or the LBD with increasing azahistidine albicidin concentrations. C) Surface model of the crystal structure of STM3175. DBD and LBD are colored in dark and light grey, respectively. The albicidin orientations of the top four best scoring models obtained with AutoDock Vina are shown in hues of blue. The inset shows cross-section through the binding pocket of the highest-ranking complex model with the conserved Phe265 highlighted in pink. The black dashed line indicates the cross-section plane. D) WebLogo (*32*) depiction of amino acids in β-strand 7 using alignments of STM3175 homologs. The conserved Phe 265 and Tyr 267 are highlighted in pink and blue, respectively (see also Fig. S18).

Attempts to crystallize the albicidin-STM3175 complex were unsuccessful but molecular docking of albicidin to the crystal structure of STM3175 suggests a binding mode where albicidin is posi-tioned in the central groove of the LBD (Fig. 2C). In the best-ranked models of the complex, the N-terminal half of albicidin occupies the binding domain groove which is a well-described binding site for other ligands of GyrI-like proteins (*31*). The C-terminal building blocks D, E and F extend through the tunnel towards the C-terminus of the LBD (Fig. 2C). Curiously, in the domain-swapped dimers found in the crystal, the exit of this tunnel is blocked by one of the helices of the DBD (Fig. S12C). However, considering the low resolution of the X-ray structure, the exact positioning of albicidin in the binding groove cannot be determined with certainty by docking studies and in some of the lower-ranked models, we also found albicidin in the opposite orientation(Figure S15A).

### STM3175 homologs in E. coli, Vibrio and Pseudomonas

A BLAST search revealed homologs in various members of the *Enterobacteriaceae* family, including *Klebsiella, Klyuvera* and *Citrobacter*. These homologs display the same transcription regulator di-domain structure with sequence identity ranging between 60 and 90%. The genomic context corresponds to that of *STM3175* in *Salmonella* with topoisomerase IV subunit A, QseB and QseC located in the direct vicinity (Fig. S16 A-B). Moreover, variants of the protein lacking the DNA-binding domain, similarly to GyrI, are present in other *Enterobacteriaceae* genera such as *Escherichia* and *Shigella*. The *E. coli* homolog is known as YgiV (approx. 50% sequence identity, similar genomic context) and a search through NCBI records revealed a number (~1500) of database entries of proteins with the same name. In the present results are the short variant (160 aa) and the di-domain protein (288 aa) with the majority of hits in *E. coli* (short variant) followed by *Salmonella enterica* and *Klebsiella pneumonia* strains (long variants).

To investigate if these homologs confer albicidin resistance and whether they are regulated by a similar GDA-based resistance mechanism, we conducted evolution experiments in *E. coli*. When exposed to increasing concentrations of albicidin, 11 out of 20 strains adapted to at least 8 μg mL^−1^ within 10 passages (Fig. 1A). Nine strains harbored GDAs, again varying in length and copy number (Fig. S17A, Table S2). A ~644 bp long region was present in all GDAs and, satisfyingly, it contained the *E. coli ygiV* gene (Fig. S15B). When expressing YgiV under control of an arabinose-inducible promoter in *S*. Typhimurium, we observed elevated albicidin tolerance comparable to that conveyed by the ligand-binding domain of STM3175 alone (Fig. S18 A-B).

AlphaFold and Rosetta both suggest that YgiV has a fold identical to that of the LBD of STM3175 with a binding groove sandwiched between two helices (AlphaFold model available in AlphaFold Protein Structure Database under UniProt ID: Q46866). Notably, the conserved Glu residue that is located at the base of the binding groove in most GyrI-like domains is also not present. In docking studies with the homology model, we obtained similar results as for STM3175 (Fig. S15B) but with a preferred orientation of the albicidin carboxylic acid towards the C-terminus of helix1 (in STM3175, this would be the end of the groove opposite of the DNA-binding domain).

We then set our sights on identifying homologs from more distant bacterial relatives. BLAST search using only the LBD of STM3175 identified single- and di-domain homologs in other *Gammaproteobacteria*, including *Vibrio sp*. and *Pseudomonas sp.*. Because of their pathogenic potential, we decided to investigate if the homologs in *Vibrio vulnificus* (YgiV-Vv; 44% identity to STM3175) and *Pseudomonas aeruginosa* (AraC-Pa-like; 48% identity) also conferred albicidin resistance. When the two di-domain homologs were expressed under control of an arabinose-inducible promoter in *S*. Typhimurium, we again observed elevated MICs, similar to those conferred by STM3175 (Figure S18 C-D).

Based on these results it is likely that single- and di-domain STM3175 homologs in various gram-negative species can increase albicidin tolerance if present in sufficient copy numbers. Most homologs (>90% of 250 BLAST search results), including YgiV-Vv and AraC-Pa, have a Phe in place of the Glu residue usually found on β-strand 7 that forms the base of the binding groove (Fig. S19). Together with a similarly conserved Tyr, this Phe residue (Phe265 and Tyr267 in STM3175) forms an ideal interaction motif for the aminobenzoic acid blocks of albicidin (Fig. 2D).

### Specificity of the resistance mechanism

To evaluate the specificity of this resistance mechanism, the six evolved *S*. Typhimurium strains (S41, T01, T04, T05, T10, T12) were challenged with compounds from different antibiotic classes, such as fluoroquinolones, tetracyclines, β-lactams or sulphonamides. With MICs comparable to those of the WT strain, none of the evolved strains showed resistance against any of the tested antibiotics (TableS6-7). Interestingly, although albicidin and fluoroquinolones both target DNA gyrase (*33*), cross resistance was not observed in either our evolved strains or fluoroquinolone resistant S*. Typhimurium* strains (FQR) (*34*) that were tested against albicidin (Fig. S20).

SbmC (GyrI) from *E*. *coli*, to which the LBD of STM3175 shows homology, protects cells from the gyrase poison microcin B17 (*35*). However, over-expression of the SbmC homolog in *Salmonella* (71% sequence identity to the *E. coli* variant), showed only a minimal increase in albicidin tolerance (Fig. S21). Likewise, despite its homology to SbmC, evolved *E. coli* strains harboring GDAs of YgiV did not convey resistance against microcin B17 (Fig. S22, TableS8). Taken together, these results indicate that the STM3175-based resistance mechanism against albicidin is highly specific.

## Discussion

In our evolution experiments, we observed the emergence of GDAs that allow *S*. Typhimurium and *E. coli* to tolerate albicidin concentrations significantly exceeding those conferred by mutations in the well-studied nucleoside transporter Tsx. This high-level resistance resulted from the presence of several copies of the transcription regulator STM3175 in the genomes of evolved strains. GDAs allow rapid transcription regulation independent of transcription factors which enables bacteria to quickly adapt to growth limiting factors (*36*) such as heat (*37*), lack of nutrients (*38*), or heavy metals (*39*).

Compared to point mutations, spontaneous duplications occur without selective pressure and at significantly higher rates (10^−4^-10^−2^/cell/division) with further increases of the copy number at rates of 10^−2^/cell/division (*3, 40*). This intrinsic genetic instability allows amplification-mediated gene expression tuning and enables populations to quickly respond to changes in environmental conditions (*36, 40*). The presence of GDAs can complicate treatment of bacterial infections as it can result in heteroresistance (*41*), in which subpopulations are less susceptible to the treatment, ultimately leading to treatment failure.

Frequently, the duplications clearly serve to increase the cellular levels of a gene product that can directly counteract the induced stress, such as efflux pumps in case of antibiotics or heavy metals (*39*). In our arabinose-inducible expression systems, the albicidin-neutralizing effect of STM3175 is clearly dosage dependent. Our adapted *Salmonella* and *E. coli* strains gain up to 15 copies of the gene and can tolerate more than a hundred-fold higher albicidin concentrations. However, GDAs are intrinsically unstable and, in the absence of selective pressure, they are generally rapidly resolved to ameliorate the inherent fitness cost (*3*). Investigations on the kinetics of *STM3175* GDA reversal is a particularly interesting aspect which will be part of future studies.

Under ongoing selective pressure, GDAs can provide a basis for the evolution of genes with new functions, as the additional gene copies could potentially evolve independently, acquiring new functions or specificities (*39, 42*), for example, to diversify or specialize an existing resistance mechanism or generation of new fusion proteins. In this study, it is conceivable that over-expression of STM3175 by GDA events paves the way for fixed, genetic albicidin resistance by subsequent mutations in the Tsx nucleoside channel, as shown for *E. coli* (*20*).

It appears unlikely that YgiV and STM3175 are resistance traits that evolved specifically to counteract albicidin. The lack of cross-resistance in our tests with numerous antibiotics and the high affinity for albicidin, however, suggest specificity of the mechanism. On the other hand, GyrI-like domain containing transcription factors like STM3175 have been implicated in polyspecific binding and multidrug resistance (*39, 42*), and STM3175 might have the ability to protect cells from other stressor molecules. A role in DNA protection has been described for the homolog GyrI. Interestingly, the protein GyrI, which was originally discovered as a gyrase inhibitor, protects DNA gyrase from the peptide toxins microcin B17 and CcdB and offers partial protection from quinolones (*31*). GyrI also imparts resistance against alkylating agents mitomycin C and N-methyl-N-nitro-N-nitrosoguanidine which act independently of gyrase (*43*).

DNA gyrase inhibitors generally fall into two categories: those that inhibit the binding of ATP and interrupt supercoiling (e.g. aminocoumarins or cyclothialidines), and those that trap enzyme–DNA intermediates (e.g. quinolones, CcdB, microcin B17). Albicidin belongs to the second category (*3–5*), hence a resistance mechanism comparable to that of microcin B17 seems reasonable. However, the molecular details behind the mechanism of action of albicidin have not yet been determined and this similarity to microcin B17 and CcdB might provide further insights.

Members of the GyrI superfamily are prevalent among bacteria, archaea and eukaryotes (*16*). PFAM lists GyrI-like family members as stand-alone domains or fused to other functional domains, such as DNA-binding or enzymatic domains. In our experiments, we observed higher resistance in over-expression experiments of full-length STM3175 compared to YgiV or STM3175 without DBD. This difference was not observed in agar diffusion experiments, which might be due to transcription regulatory activity of the protein. It would be interesting to see in future experiments which promoter regions are recognized by the DBD and which genes are directly affected.

A number of structures of proteins with GyrI-like domains are available in the PDB but only a few of the deposited structures are of di-domain proteins: ROB (cite Kwon et al. doi: 10.1038/75213.), BmrR and EcmrR, two multi-drug sensing regulators of the MerR family that consist of a N-terminal helix-turn-helix DNA-binding domain with a dimerization motif and a GyrI-type ligand binding domain (cite: Bachas et al 2011, doi: 10.1073/pnas.1104850108 and Yang et al 20211, doi: 10.1038/s41467-021-22990-8). The domain architecture of STM3175 resembles that of ROB which also consists of an N-terminal AraC DNA-binding domain fused to a GyrI-like domain (cite Kwon et al. doi: 10.1038/75213). Despite its relatively low resolution, the crystal structure of STM3175 agreed very well with predictions and available GyrI-family structures in the PDB. And while we were not able to obtain crystals of albicidin bound to STM3175, our models suggest a binding mode that is consistent with structural data of various drug-like compounds in complex with GyrI-like proteins (Moreno et al and Yang et al and Bachas et al).

In summary, we demonstrated that exposure of *S*. Typhimurium and *E. coli* to increasing concentrations of the gyrase poison albicidin results in rapid adaption via chromosomal duplication-amplification events. The affected region harbors the GyrI-like domain containing transcription regulator STM3175 (YgiV), which we identified as a critical factor involved in high-level resistance against albicidin. The protein binds albicidin with high affinity in an equimolar stoichiometry. We further showed that this resistance mechanism/gyrase protection mechanism is ubiquitous in *Enterobacteriacea* with STM3175 homologs conferring resistance in *Escherichia, Vibrio* and *Pseudomonas*.

## Supporting information

PDB report

## Acknowledgments

The FP7 WeNMR (project# 261572), H2020 West-Life (project# 675858), the EOSC-hub (project# 777536) and the EGI-ACE (project# 101017567) European e-Infrastructure projects are acknowledged for the use of their web portals, which make use of the EGI infrastructure with the dedicated support of CESNET-MCC, INFN-PADOVA-STACK, INFN-LNL-2, NCG-INGRID-PT, TW-NCHC, CESGA, IFCA-LCG2, UA-BITP, SURFsara and NIKHEF, and the additional support of the national GRID Initiatives of Belgium, France, Italy, Germany, the Netherlands, Poland, Portugal, Spain, UK, Taiwan and the US Open Science Grid. Stefan Schwarz (Institute of Microbiology and Epizootics, Freie Universität Berlin) is kindly acknowledged for providing strains FQR1-4 and we are thankful to Peter Schwerk and Julian Brombach (Institute of Microbiology and Epizootics, Freie Universität Berlin) for excellent technical assistance. We accessed beamlines of the BESSY II (Berliner Elektronenspeicherring-Gesellschaft für Synchrotronstrahlung II) storage ring (Berlin, Germany) via the Joint Berlin MX-Laboratory sponsored by the Helmholtz Zentrum Berlin für Materialien und Energie, the Freie Universität Berlin, the Humboldt-Universität zu Berlin, the Max-Delbrück-Centrum, the Leibniz-Institut für Molekulare Pharmakologie and Charité – Universitätsmedizin Berlin. We are grateful to Elvenstar’s Pukipon for the kind donation of cat whiskers. For mass spectrometry, we would like to acknowledge the assistance of the Core Facility BioSupraMol supported by the Deutsche Forschungsgemeinschaft (DFG).

## Author contributions

MF, SK, MSa, KT, RS conceived the experiments, SK, MSa, MSe ND, BL, WC, SW, JK, FG, BK, KT, DG, JGH, SG and TS performed the experiments and analyzed the data; SK and MF wrote the paper. All authors discussed the results and contributed to the final manuscript.

## Data and materials availability

The coordinates of the X-ray crystal structure have been deposited with the Protein Data Bank, Research Collaboratory for Structural Bioinformatics at Rutgers University (PDB-ID: 7R3W). All data is available in the main text, the supplementary materials or, upon request, from the authors. The mass spectrometry proteomics data have been deposited to the ProteomeXchange Consortium via the PRIDE partner repository with the dataset identifier PXD031944.

## Supplementary Materials

Materials and Methods

Figures S1-S22

Tables S1-S13

References (1 - 34)

