## Supplemental Files for "Gene amplifications cause high-level resistance against albicidin in Gram-negative bacteria"

####Supplementary Materials for

### Gene amplification of a GyrI-like transcriptional regulator confers high-level resistance against albicidin in Gram-negative bacteria

**Authors:** Mareike Saathoff[1#], Simone Kosol[2#], Torsten Semmler[3], Karsten Tedin[1], Nicole Dimos[4], Johannes Kupke[1], Maria Seidel[2], Fereshteh Ghazisaeedi[1], Silver A. Wolf[3], Benno Kuropka[4], Wojciech Czyszczoń[5], Dmitry Ghilarov[5], Stefan Grätz[2], Jonathan G. Heddle[5], Bernhard Loll[4], Roderich D. Süssmuth[2#] and Marcus Fulde[1,6#]

**This PDF file includes:**

Materials and Methods
Figs. S1 to S22
Tables S1 to S13

#### Materials and Methods

*Bacteria, media and antibiotics*

The isolates and plasmids used in this study are described in Table S9. LB Lennox (Roth, Karlsruhe, Germany) was used as broth or in agar plates for all experiments. Incubations steps took place at 37°C and 200 rpm. Throughout the study, azahistidine albicidin was used due to its increased stability and similar activity compared to natural albicidin (*1*) (Fig. 1). If treatment with the antibiotic albicidin was required, incubation followed under light exclusion. When necessary antibiotics were supplemented with the following concentrations: carbenecillin (100 µg $mL^{-1}$; Roth, Karlsruhe, Germany), chloramphenicol (15 µg $mL^{-1}$; Sigma-Aldrich, Taufkirchen, Germany), kanamycin (50 µg $mL^{-1}$; Roth, Karlsruhe, Germany).

*Antimicrobial susceptibility testing*

Minimum inhibitory concentrations (MIC) assays were performed according to Clinical and Laboratory Standards Institute (CLSI) recommendations in LB Lennox broth in flat-bottomed, 96-well plates (Corning, Wiesbaden, Germany) with an inoculum of $10^6$ bacteria/well and incubated overnight. The optical density of the cultures was determined at $\lambda$ = 460 and 600 nm immediately after inoculation and after overnight incubation using a BioTek Synergy HT plate reader. A serial dilution of albicidin or Microcin B17 in 100 % dimethylsulfoxid (DMSO; Sigma-Aldrich, Taufkirchen, Germany) was necessary to ensure a constant DMSO concentration of 5 % (v/v). Five µl of each dilution step was pipetted into a 96-well plate. Afterwards 95 µl of the prepared cells were added to the albicidin serial dilution. Controls included medium alone and wells without antibiotic additions with 5 % DMSO and without DMSO. MIC assays with ciprofloxacin were performed with an inoculum of $10^5$ bacteria/well. Antimicrobial susceptibility testing was performed by broth micro-dilution for evolved and control strains according to the guidelines of the Clinical and Laboratory Standards Institute. Sensititre test plates NLD1VMON, NLD2VMON and NLD3VMON (Thermo Fisher Scientific, Schwerte, Germany) were used.

For agar diffusion assays with microcin B17 bacteria were adjusted to McFarland Standard 0.5 and streaked out on LB Lennox agar plates. Ten µl of 10 mg $ml^{-1}$ microcin B17 were pipetted to a blank filter in the middle of the plate, incubated at 37°C and the zone of inhibition was detected after 24 h.

Experiments were basically performed with three biological and three technical replicates. Any differences were stated in the captions of the respective figure.

*Evolution experiments with albicidin*

For the evolution experiment 6 mL LB broth containing 0.06 µg $mL^{-1}$ azahistidine albicidin were inoculated with 10 µl of an overnight culture of the wild type strain *Salmonella enterica* subsp. *enterica* serovar Typhimurium ATCC 14028 After 24 h of incubation resistance was determined by visible growth (OD measurement). The evolution procedure was performed in 3 independent experiments (N=30 each) staring with one ancestral strain.

For evolution with the *Δtsx* mutant strain (9866) starting with one ancestral strain 10 independent experiments were performed. Therefore, 6 ml starter culture was inoculated with 10 µl of an overnight culture. The albicidin concentration was stepwise enhanced within nine passages beginning with the half MIC (0.125, 0.25, 0.5, 1, 1.5, 2, 4, 8, 16 and 20 µg $mL^{-1}$) of the input strain, whereby the transfer volume ranged from 5 to 20 µl depending on the detected turbidity. To ensure a stable resistance phenotype all positive samples were treated a second time with albicidin.

Evolution experiments with wild type strain *E. coli* ATCC 25922 were started with 1-fold MIC (0.0156 µg $mL^{-1}$) and passaged 10 times with increasing albicidin concentrations to a maximum of 8 µg $mL^{-1}$ in 20 independent approaches.

*Cloning and overexpression*

For overexpression in an arabinose-inducible expression vector, the genes *STM3175* (ACY90250.1), *ygiW* (ACY90251.1) and *qseB* (ACY90252.1) were PCR amplified using primers 3175XbaIF/3175Hind3R (*STM3175*), YGIWXbaIF/YGIWHind3R (*ygiW*) and PREAEcoRI/PREAXbaIR (*qseB*). *Sbmc* from *S.* Typhimurium was amplified with the primer pair SBMCXbaIF/ SBMCHind3R. *STM3175*, *ygiW*, *ygiV* and *sbmC* from *S.* Typhimurium were digested with *XbaI*/*HindIII*, whereas *qseB* was digested with *EcorI*/*XBaI*. All Fragments were inserted into plasmid pBAD30 (see Table S8). *STM3175-DBD* and *STM3175-LBD* were amplified by inverse PCR using high fidelity Taq polymerase with pBAD30_STM3175 as template (Primer: 3175XhoIF/3175XhoIR (*STM3175-LBD*), 3175-Xho-AraC-F2/3175-Xho-AraC-R (*STM3175-DBD*). PCR constructs were *XhoI* digested, re-ligated and inserted into plasmid pBAD30.

Furthermore, *ygiV* and *ygiW* from *E. coli* were cloned as a *XbaI*/*HindIII* PCR fragment with primer pair ECygiVXF/ECygiVH3R into pBAD30 and *E. coli ygiW* was amplified with primer pair ECygiWXF/ ECygiWH3R.

For overexpression of *ygiV* from *V. vulnificus* (*ygiV*-Vv) and *araC* from *P. aeruginosa* (*araC*-Pa), genes were designed and ordered from GenScript and delivered in plasmid pUC57. Genes were amplified with primer pUC/M13(-40)/ pUCM13/Rev and digestion followed for *ygiV*-Vv with KpnI/SalI and *araC*-Pa with EcoRI/BamHI. Both genes were inserted into pBAD30. S*almonella enterica* subsp. *enterica* serovar Typhimurium

ATCC 14028 was transformed with the plasmids (pBAD30-STM3175, pBAD30-STM3175-DBD, pBAD30-STM3175-LBD, pBAD30-ygiW, pBAD30-qseB, pBAD30-sbmC, pBAD30-ygiV (*E. coli*) and pBAD30-ygiW (*E. coli*), pBAD30-ygiv (*V. vulnificus*) and pBAD30-araC (*P. aeruginosa*), see Table S9. Overexpression was induced with at least 1 mM arabinose.
For overexpression of STM3175 in an N-terminal His-tagged T7 vector (pET28a(+)) several STM3175-constructs were generated (see Table S9) by PCR amplification from template pBAD30_STM3175 with different primer combinations including a TEV-site (GAAAACCTGTATTTTCAGGGC) for later truncation and further protein analyses (see Table S10). PCR-products were digested with *NdeI*/*BamHI* and inserted into pET28a(+). All STM3175 gene-constructs were expressed in *E. coli* BL21-Gold (λDE3) with the T7 promotor expression system including an N-terminal TEV-cleavable His-tag. Expression was induced with 0.1 mM isopropyl β-D-1-thiogalactopyranoside (IPTG). The expression culture was further incubated for 20 h, 200 rpm at 20 °C. YgiV (STM3175 homolog) from *E. coli* was cloned as Ndel/HindIII PCR fragment with primer ECygiVNF and ECygiVH3R into pet28a(+). Primers were purchased from Invitrogen (Karlsruhe, Germany) and enzymes were obtained from Promega (Walldorf, Germany).

*Sequencing and data analysis*
Sequencing of the *tsx* genes were performed by PCR isolation (QiAquick PCR Purification Kit, Qiagen) and further sequenced with primer TsxF/TsxR by LGC genomics. For whole genome sequencing DNA was isolated with the QIAamp DNA Mini Kit (Qiagen).
Short-read sequencing was performed on an Illumina NextSeq 2000 sequencer with NextSeq 2000 P1 Reagents (300 Cycles) (Illumina Inc., San Diego, CA, USA), and the Library Preparation Kit Nextera XT (Illumina), resulting in 150 bp paired-end reads and a ca. 100-fold coverage on average. Additionally, long-read sequencing was performed using Oxford Nanopore MinION (Oxford, UK). MinION one-dimensional (1D) libraries were constructed, using the SQK-RBK004 kit (Nanopore Technologies, Oxford, UK) and loaded according to the manufacturer's instructions onto an R9.4 flow cell. The sequencing data was collected for 48 h. Total amounts of 1 ng and 400 ng extracted DNA were used as starting material for sequencing by Illumina NextSeq and MinION, respectively. A closed genome was generated by a de-novo hybrid assembly using a combination of short and long reads with Unicycler v0.4.7 (*2*).

*RNAseq*
The wild type and mutant T12 *Salmonella* Typhimurium strain were streaked out on LB Lennox agar plates, incubated at 37°C overnight and 6 ml LB Lennox broth were inoculated with a single colony. After overnight incubation at 37°C, 200 rpm three sterile LB-Lennox broth cultures were inoculated with 50 µL of the overnight culture and cultivated until an $OD_{600nm}$ = 0.5. RNA was isolated with RNeasy Mini Kit (Qiagen) according to instructor's manual of the RNAprotect Bacteria reagent handbook (protocol 4: Enzymatic Lysis and Proteinase K Digestion of Bacteria and protocol 7: Purification of Total RNA from Bacterial Lysate using the RNeasy Mini Kit). The experimental setup includes three biological replicates, each performed in triplicates.
RNA sequencing was performed on an Illumina MiSeq using 250bp paired-end reads resulting in samples of 4,199,984 (8055) 5,647,216 (8054) and 5,146,530 (8053) reads, respectively. Reads were adapter-trimmed and quality-filtered using fastp v0.23.2 with default settings. Next, trimmed reads were imported into the Geneious Prime v2020.2.3 software and mapped to the corresponding reference genome using Geneious assembler with default settings. Transcript quantification was performed from within Geneious based on annotated CDS using the "count as partial matches" setting for ambiguous reads. Expression level metrices (including log2 ratio, TPM and counts) were subsequently calculated by comparing the mapped reads between 8053 and 8055, and were exported as Excel tables. Fold changes were visualized in Geneious and heatmaps were generated using the ComplexHeatmap v2.8.0 package in R v4.1.2.

*Proteomics*

*Growth conditions and protein extraction*
The wild type (WT) and evolved (T12) IMT-9866 *Salmonella* Typhimurium strains were cultured on LB-agar at 37°C overnight. Subcultures of strains with an $OD_{600}$ of 0.5 were prepared by inoculating 10 ml of LB broth with a single colony (five replicates per strain). Samples were centrifuged for 10 min at 4°C at 10000 × g. The pellets were resuspended in 80% ethanol (v/v) and the centrifugation step was repeated. Air dried bacterial pellets were resuspended in 200 µl of 20 mM ice-cold HEPES buffer. Cell disruption was performed by sonication (UP100H; Hielscher Ultrasound Technology, Teltow, Germany) using a duty cycle of 1.0 and an amplitude of 100% for 45 sec on ice. Samples were centrifuged for 10 min at 4°C at 12000 × g and supernatants were collected. Protein quantification was done by the Qubit Protein Assay and quality control of protein extraction was done by SDS-PAGE (*3*).

*In-solution trypsin digestion*

Five µg of total extracted protein per sample were acetone precipitated. The pellet was reconstituted in 20 µL of denaturation buffer (6 M urea/2 M thiourea in 10 mM HEPES, pH 8.0). The next steps were carried out following the protocol from Wareth et al. (*4*). In brief, 0.2 µl of 10 mM DTT was added to each sample and reduction was carried out for 30 min at room temperature (RT). Then 2 µl of 55 mM iodoacetamide was added and alkylation was carried out for 20 min at RT in the dark. Subsequently, 0.1 µg of LysC (125-05061 Wako Chemicals) was added and protein digestion was performed overnight at RT. The next day, samples were diluted with 50 mM ammonium bicarbonate (1:4) to reduce the urea concentration below 2 M. Finally, 0.1 µg of trypsin was added, and protein digestion was carried out overnight at RT. The digestion was stopped by acidifying the samples to pH <2.5 by adding 100 µl of 5% acetonitrile (ACN) and 3% trifluoroacetic acid (TFA) in water. Following the digestion, peptide samples were desalted by solid phase extraction (SPE) using C18 stage tips (*5*).

*Nano liquid chromatography-mass spectrometry (LC-MS) and data analysis*

Dried peptides were reconstituted in 30 µl of 0.05% TFA, 4% ACN, and 3 µl were analyzed by an Ultimate 3000 reversed-phase capillary nano liquid chromatography system connected to a Q Exactive HF mass spectrometer (Thermo Fisher Scientific). First, peptides were injected and concentrated on a trap column (PepMap100 C18, 3 µm, 100 Å, 75 µm i.d. x 2 cm, Thermo Fisher Scientific) equilibrated with 0.05% TFA in water. After switching the trap column inline, LC separations were performed on a capillary column (Acclaim PepMap100 C18, 2 µm, 100 Å, 75 µm i.d. x 25 cm, Thermo Fisher Scientific) at an eluent flow rate of 300 nl/min. Mobile phase A contained 0.1 % formic acid in water, and mobile phase B contained 0.1% formic acid in 80% ACN / 20% water. The column was pre-equilibrated with 5% mobile phase B followed by an increase of 5–44% mobile phase B in 70 min. Mass spectra were acquired in a data-dependent mode utilising a single MS survey scan (m/z 350–1650) with a resolution of 60,000, and MS/MS scans of the 15 most intense precursor ions with a resolution of 15,000. The dynamic exclusion time was set to 20 seconds and automatic gain control was set to $3x10^6$ and $1x10^5$ for MS and MS/MS scans, respectively.

MS and MS/MS raw data were processed and analyzed using the MaxQuant software package version 1.6.14 with implemented Andromeda peptide search engine (*6*). Data were searched against the custom database for *Salmonella* Typhimurium with 5000 protein sequences generated from the whole genome sequence of IMT-9866 *Salmonella* Typhimurium strain from our lab. The default settings of MaxQuant were used except for enabling label-free quantification (LFQ) and match between runs (MBR). Filtering and statistical analysis was carried out using the Perseus software version 1.6.14 (*7*). After initial filtering (removing contaminants, reverse hits and hits only identified by site), only proteins which were quantified with LFQ intensity values in at least 3 replicates (within at least one of the two experimental groups) were used for downstream analysis. Missing values were replaced from a normal distribution (imputation) near the detection limit using the default settings (width 0.3, down shift 1.8). Mean $\log_2$ fold protein LFQ intensity differences between the 2 experimental groups (background strain - evolved strain) were calculated in Perseus using student's t-test with a permutation-based FDR of 0.05. The volcano plot was created by plotting the $-\log_{10}$ p-values against the mean $\log_2$ fold protein LFQ intensity differences. Proteins were considered significantly changed between experimental groups if they have a q-value < 0.05 and at least a 2-fold change in LFQ intensity ($\log_2$ fold change >1 for the evolved strain or $\log_2$ fold change <-1 for the background strain, respectively). The mean log2 fold protein LFQ intensity differences of GDA region proteins are displayed in a heat map visualization using GraphPad Prism version 9 for Windows, GraphPad Software, San Diego, California USA, www.graphpad.com. All identified and quantified proteins including the volcano plot are shown in Table S13.

*Bioinformatics – Sequence Analysis and homology modelling*

To identify the characteristic read coverage of the GDA region, Illumina sequencing reads of the evolved strains were trimmed and mapped to the closed genome of the respective WT strain using QIAGEN CLC Genomics Workbench v20 (Qiagen) with standard parameters, linear gap costs. Non-specific matches were mapped randomly.

To identify conserved domains and protein homologs, the amino acid sequence of STM3175 from *Salmonella* Typhimurium was submitted to a Basic Local Alignment Search Tool (BLAST) search including conserved domain search (CCD) (*8*). To search specifically for homologs of the ligand-binding domain in other bacterial genera, the BLAST search was repeated using only the sequence of that domain as input, excluding Salmonella from the results. Multiple sequence alignments were generated with Clustal Omega (*9*).

The PSIPRED Server (*10*) was used for secondary structure predictions of AlbA and STM3175. The molecular structures of STM3175 from *S.* Typhimurium and ygiV from *E. coli* were modelled using the Protein Fold Recognition Server PHYRE2 (*12*) and the RoseTTAFold method of Robetta (*13*). Homology models were assessed in pymol and UCSF ChimeraX (*11*).

*Molecular Docking*

Albicidin was docked with a monomer of the X-ray crystal structure of STM3175 using AutoDock Vina (*14*) in UCSF ChimeraX. The homology models of STM3175 and YgiV were docked with DockingVina or HADDOCK (*15*). The Robetta homology model of full-length STM3175 was used for docking in HADDOCK. To compare the docked complexes of both proteins, STM3175 and YgiV, the structures modelled by PHYRE2 using SbmC as template were submitted to the programs. Albicidin was introduced in a starting conformation resembling its bound state to AlbA. Based on published recognition site data for the GyrI-like superfamily (*16*), the binding groove and hydrophobic amino acids along the central groove located between the two SH2 domains were chosen as active residues and docking was carried out using standard parameters.

*Agar diffusion assay*

An overnight culture of *E. coli* DSM1116 was diluted to an $OD_{600}$ of 0.05 in LB agar (10 g/L peptone, 5 g/L yeast extract, 5 g/L NaCl, 0.75% agar agar, pH 7.85). 20 mL of the inoculum were poured into sterile petri dishes and cooled down to room temperature for 30 minutes. Subsequently, holes with a diameter of 5 mm were punched into the agar plates with the end of a glass pipette. The protein-albicidin reaction mixtures contained 37.5 µM azahistidine-albicidin and 37.5 µM (1:1) or 18.75 µM (2:1) purified protein STM3175 or LBD, respectively. The positive controls consisted of only 37.5 µM azahistidine-albicidin in 5% dimethyl sulfoxide (DMSO)/ assay buffer (50 mM Tris, 200 mM NaCl, 1 mM EDTA, pH 8.5). The negative control consisted of 37.5 µM (1:2) or 18.75 µM (1:1) purified protein STM3175 or LBD in 5% DMSO/assay buffer, respectively. Following incubation (30 min, RT, in the dark), 30 µL of each reaction mixture and controls were transferred into the holes of the agar plates. Plates were dried for 30 min and subsequently incubated at 37°C overnight.

*Purification of recombinant STM3175 and STM3175-LBD*

*E. coli* BL21(DE3) cells were transformed with pet28a (+)-STM3175 or pet28-STM3175-LBD, respectively. LB broth supplemented with 50 µg/ml kanamycin was inoculated from over-night cultures and grown at 37°C until an $OD_{600}$ of 0.6 was reached. After induction with 0.1 mM isopropyl β-D-1-thiogalactopyranoside (IPTG) the proteins were expressed at 18°C for 20 hrs. The cells were harvested by centrifugation, resuspended in lysis buffer (50 mM Tris, pH 8.5, 500 mM NaCl, 10 mM glucose, 10% glycerol) and processed in a cell disruptor (Constant Systems Ltd) at 25 kPsi before centrifugation at 50 000 x G for 45 min. The lysate was incubated with Pure Cube Ni-NTA agarose (Cube Biotech, Germany) at 4°C for 20 min while shaking and after a washing step with lysis buffer, the protein was eluted with 250 mM imidazole in lysis buffer. $His_6$-STM3175 and $His_6$-STM3175-LBD were buffer exchanged in gel filtration buffer (50 mM Tris, pH 8.5, 200 mM NaCl, 1mM EDTA) on PD10 Desalting Columns and the purity of the proteins was assessed on SDS-PAGE (Fig. S10A). For crystallization experiments and CD spectroscopy, the proteins were further purified on a Superdex HiLoad 16/60 g75 gel filtration column (GE Healthcare). Before gel filtration, $His_6$-STM3175-LBD was cleaved over night with GFP-TEV (1:10 molar ratio) and separated from GFP-TEV and the His-tag peptide using a 1ml HisTrap HP column (GE Healthcare). Gel filtration fractions containing $His_6$-STM3175 or STM3175-LBD were pooled and concentrated to 20 mg/ml. Protein concentrations were determined on a nano-photometer P330 (Implen, Munich, Germany) using the extinction coefficient of 60055 $M^{-1}$ $cm^{-1}$ as calculated by the ProtParam tool (Expasy)..

*CD spectroscopy*

STM3175 and STM3175-LBD were dissolved in buffer (50 mM Tris, pH 8.5, 200 mM NaCl, 1mM EDTA) at concentrations of 7.5 µM and 10 µM, respectively. CD spectra were acquired at 20°C on a JASCO with a bandwidth of 1.0 nm and a scanning speed of 5 nm/min in 5 accumulations. The data pitch was set to 0.1 nm. The spectra were processed using spectra analysis software

$$MRE = \frac{\theta * 0.1}{l * c * n} \qquad (1)$$

Where $\theta$ is the ellipticity and $l$, $c$, and $n$ denote the path length, molar concentration and number of amino acids.

*Tryptophan fluorescence quenching*

For tryptophan fluorescence quenching measurements, azahistidine albicidin stock solutions of 1200, 600, 200, 100, 50, 25, 12.5 and 6.25 µM were prepared in assay buffer (50 mM Tris, pH 8.5, 200 mM NaCl, 1mM EDTA) with 20% DMSO. The prepared solutions were added to 200 nM STM3175 or 600 nM STM3175-LBD in assay buffer to yield final concentrations of 6, 3, 1, 0.5, 0.25, 0.125, 0.062 and 0.031 µM azahistidine albicidin. Before spectra acquisition, the samples were incubated at room temperature for 15 min in the dark. Tryptophan fluorescence was monitored on a FluoroMax 2 spectrometer from Horiba (Potsdam, Germany) with the excitation wave length set to 280 nm. Emission spectra were recorded from 290–450 nm with a scanning speed of 60 nm $s^{-1}$ and an integration time of 1 s. The excitation and emission slit widths were set to 5 nm. All measurements were performed in triplicate and standard deviations are given for the $K_d$ and Hill factor n. The quenching factor was determined from the normalized integrated emission band (290-400 nm) of each titration step subtracted from the emission band of the protein without ligand. The normalized tryptophan fluorescence quenching data was fitted to the equation 2:

$$Y = B_{max}\frac{[L]^n}{K_d^n+[L]^n} \qquad (2)$$

where [L] is the total ligand concentration, n is the Hill coefficient, and $B_{max}$ denotes the maximum binding capacity of the protein.

*Microcin B17 purification*

*E. coli* BW25113 cells were transformed with a plasmid carrying the whole microcin B17 (MccB17) operon (*pBAD-mcbABCDEFG*) (*17*) required for biosynthesis. Cells were grown at 37°C in LB media until an $OD_{600}$ of 0.6 was reached before induction with 1 mM arabinose and expression of MccB17 overnight at 30°C. Cells were harvested (5000 x G for 15 min), resuspended in 100 mM acetic acid/1 mM EDTA solution and lysed by boiling in a water bath for 10 min. The lysate was cleared by centrifugation (12 000 x G for 30 min) and filtered with a 0.22 µm on-syringe filter before loading onto a C18 cartridge (Agilent BondElut, HF Mega BE-C18) equilibrated with 0.1% TFA. After loading, the cartridge was washed with 10% MeCN in 0.1% TFA, then bound MccB17 was eluted with 30% MeCN in 0.1%TFA in 10 ml fractions and freeze-dried overnight. The crude MccB17 pellet was subsequently redissolved in 100% DMSO and purified by HPLC (COSMOSIL 5C18-MS-II Packed column, 120Å, 5 µm, 10.0 mm ID x 150 mm). MccB17 was loaded onto the 0.1%TFA pre-equilibrated column in portions of 450 µl of 10% MccB17-DMSO in 0.1%TFA and eluted by a 10-30% gradient of MeCN in 0.1%TFA. MccB17 eluted after 14 and 17 min, individual peaks corresponding to fully matured forms of MccB17 were collected and freeze-dried.

*Crystallographic methods*

For crystallization experiments STM3175 was concentrated to 18.4 mg/ml. Initial crystals were obtained by the sitting-drop vapor-diffusion method at 4 °C with a reservoir solution containing 0.1 M magnesium formate. Intergrown crystals were used to prepare a seed stock. With a cat whisker, seeds were transferred to a freshly prepared crystallization drop. Well-formed crystals were soaked in 30% glycerol plus reservoir solution and frozen in liquid nitrogen. Data was collected at Berlin BESSYII, beamline 14.2. X-ray data collection was performed at 100 K. Diffraction data were processed with the XDS (*18*) in space group $C222_1$ (Table S12). The structure of the STM3175 was solved by molecular replacement. A search model of full-length STM3175 was prepared with ROBETTA (*13*). An initial search with full-length STM as search model using PHASER (*19*) failed. Next, the model was divided in a N- and C-terminal corresponding approximately to half of the full-length protein. PHASER could locate 4 copies of the C-terminal domain. Subsequently, a search with the N-terminal fragment, revealed only one copy. Careful inspection of the electron density indicated the presence of further secondary structure elements, which were manually placed and extended which finally allowed to place a total four copies of the N-terminal domain. The structure was refined by maximum-likelihood restrained refinement using PHENIX (*20, 21*) followed by iterative, manual model building cycles with COOT (*22*). Model quality was evaluated with MolProbity (*23*). Figures were prepared using PyMOL (Schröderinger Inc.).

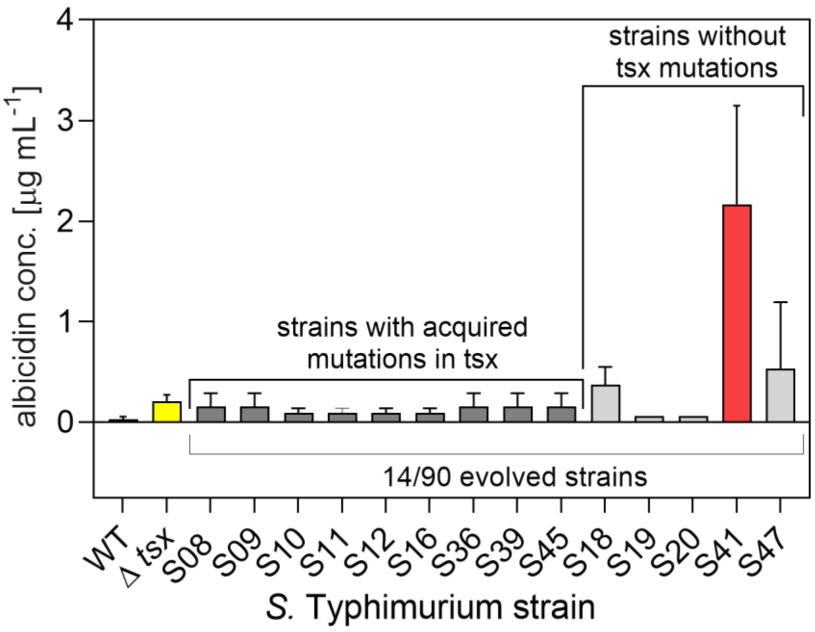

**Fig. S1. MIC detection of evolved *S.* Typhimurium strains.** Adaptation to albicidin in 14/90 strains after overnight incubation with the 4-fold albicidin wild type MIC (WT; ATCC 14028; black bar). Nine evolved strains (dark grey bars) showed mutations in the *tsx* gene and a comparable MIC to the *tsx* mutant (white bar, 0.25 µg $mL^{-1}$). Five evolved strains showed 100% *tsx* gene identity to the WT. The strain S41 (red bar) has an increased MIC of 2 µg $mL^{-1}$, corresponding to an increase by almost 70-fold to the WT (0.00156 µg $mL^{-1}$). This strain was investigated in further experiments. Data represent mean and standard deviations of six biological replicates that were performed in three technical replicates (WT, Δ*tsx*, S41). The data of the other strains represent means and standard deviations of two biological replicates that were performed in three technical replicates. Strains without error bar have the same result in every single experiment.

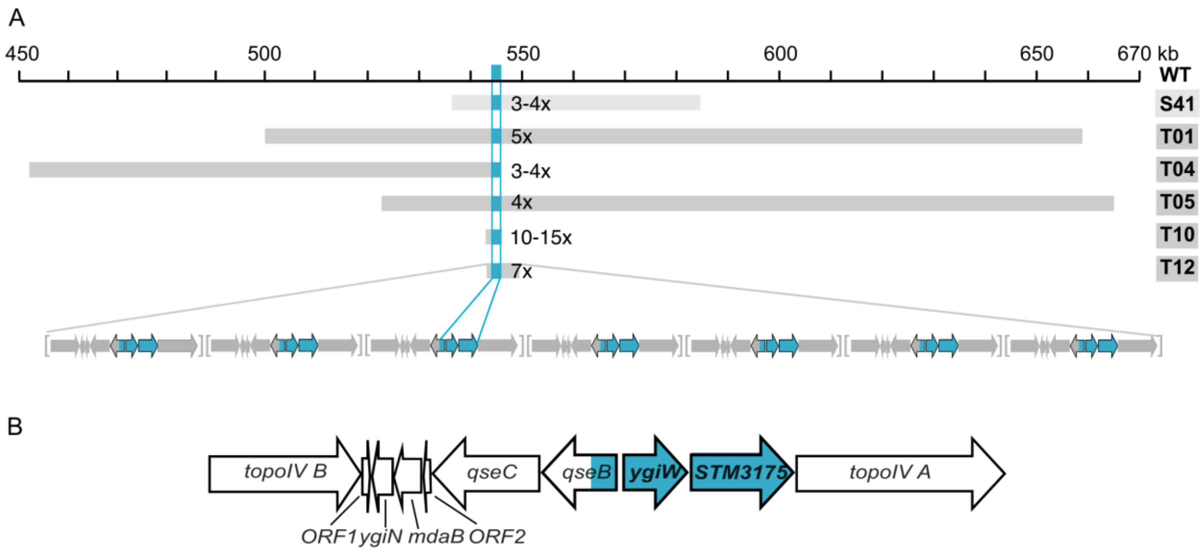

**Fig. S2. Albicidin high-level resistance resulting from GDAs in evolved *S.* Typhimurium.** A) Evolution of GDAs in six strains after treatment with albicidin (S41: 0.06 µg $mL^{-1}$ albicidin for 24 h, light grey bar; T-strains: increasing albicidin conc. from 0.125 to 20 µg $mL^{-1}$ in nine passages, dark grey bars) results in multiple copy regions of varying length. The same ~2200 bp segment is present in each copy region in all evolved strains (cyan). T12 with seven copies of a region is shown as example. B) The amplified region in T12 contains the genes for topoisomerase IV subunits A and B (*topoIV A*, *topoIV B*), QseBC, STM3175, YgiW, YgiN, modulator of drug activity B (*mdaB*) and two ORFs for hypothetical proteins.

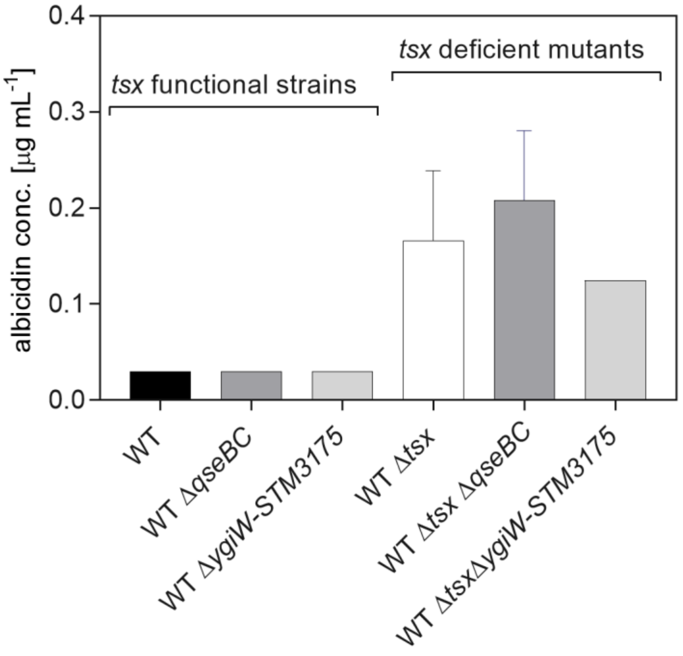

**Fig. S3. MIC determination of knockout mutants.** *S*. Typhimurium knockout mutants of *qseBC* (dark grey) and *STM3175-ygiW* (light grey) show the same azahistidine albicidin tolerance as their respective parent strain: WT (black) or Δ*tsx* mutant (white). The data represent means and standard deviations of three biological replicates that were performed in three technical replicates. Strains without error bar have the same result in every single experiment.

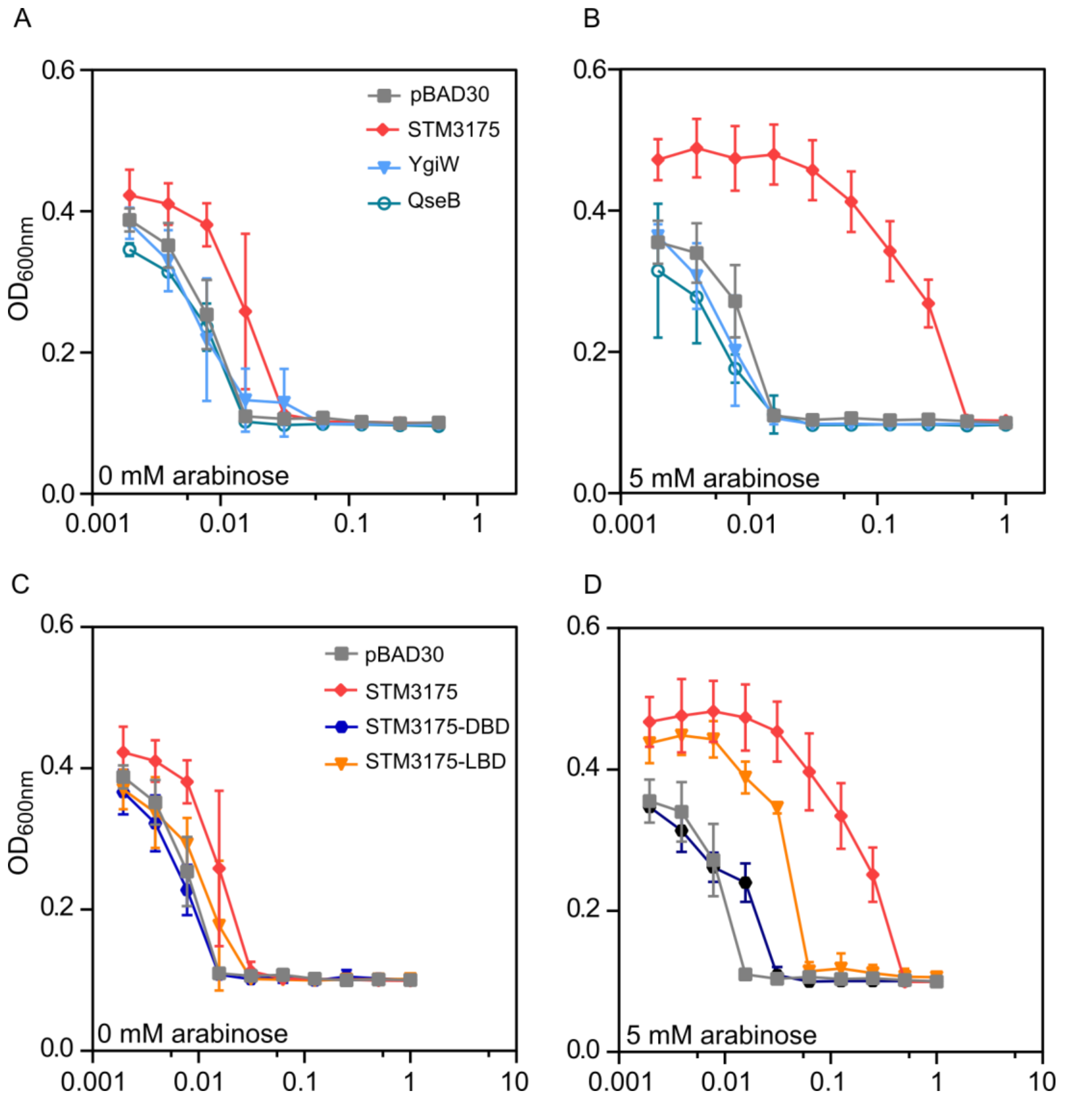

**Fig. S4. MIC curves of the STM3175 operon and domains.** A-B) Comparison of non-induced and arabinose induced expression systems of genes in the GDA region shows increased azahistidine albicidin tolerance when expression of STM3175 is induced. C-D) After supplement of 5 mM arabinose the MIC was increased for full length STM3175 and, to a lesser extent, its LBD. The data represent means and standard deviations of three biological replicates that were performed in three technical replicates

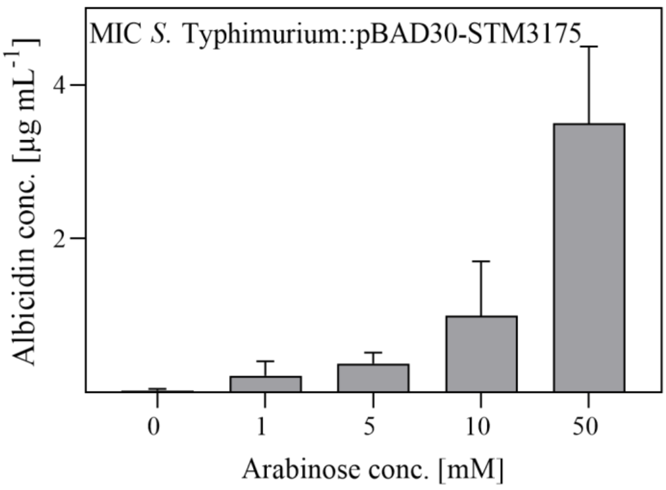

**Fig. S5**. **MIC determination for *S*. Typhimurium with STM3175-overexpression**. Overexpression of STM3175 was induced by increasing arabinose concentrations. The data represent means and standard deviations of four biological replicates that were performed in three technical replicates.

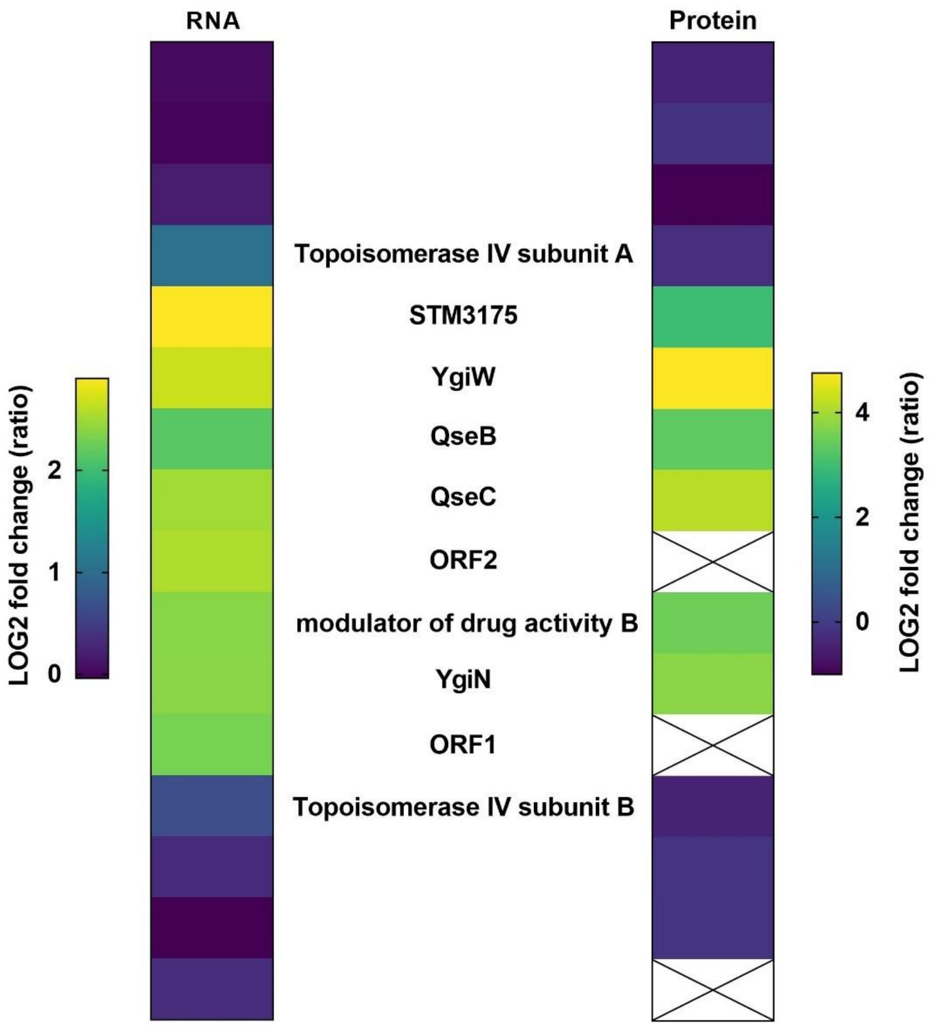

**Fig. S6. Upregulation of the GDA region.** RNA sequencing and proteomics data of evolved *S*. Typhimurium strain T12 show elevated mRNA and protein expression levels for genes in the GDA, particularly STM3175 and YgiW. Proteins indicated by a cross were not identified in the proteomics analysis.

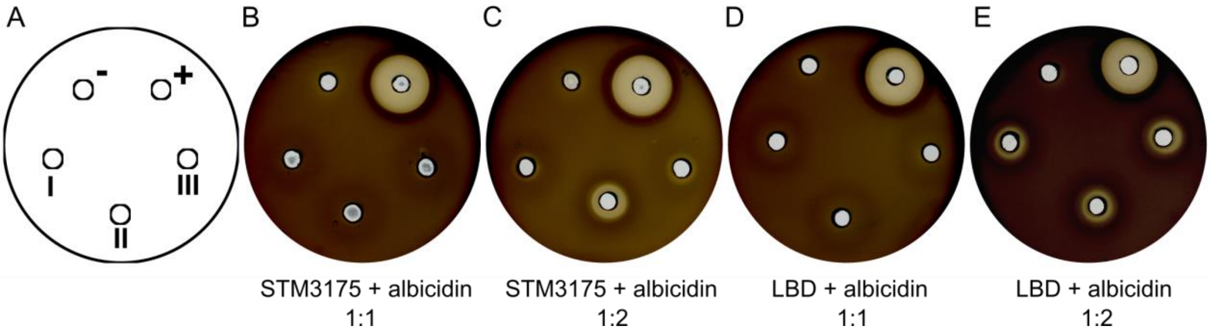

**Fig. S7. Agar diffusion assays**. STM3175 and LBD (GyrI-like domain) with azahistidine albicidin. A) Assay scheme illustrating the sample arrangement on agar plates. The negative and positive controls contain only protein (-) or azahistidine albicidin (+) in buffer with 5% DMSO, respectively. B-E) STM3175 and LBD with albicidin in a 1:1 or 1:2 molar ratio (in triplicates I-III).

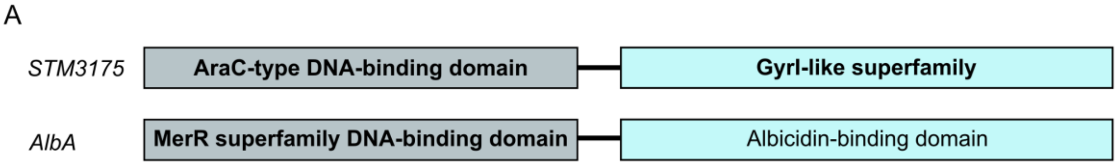

B

```
STM3175 Salmonella     ------------------------------------------------------------ 0
AlbA Klebsiella        MLEMLIQVGELAKRAGMTVRTLHHYEQTGLLTPSARSAAGYRLYNLAAVQRLHMIKALAQ 60

STM3175 Salmonella     -------MNDLIS-AA--YSERLRRVCDHIERHLDEPLSI--------EALSRM------ 36
AlbA Klebsiella        AGLELATIKDYLDRDAFSLSDLLVKQIATLDRQLQTLSTLRQRLALLREELDSGGEPDLE 120
                              ::* :.  *   *: * :    ::*:*:   ::        * *.

STM3175 Salmonella    AHSSPFHFHRQFTTWSGLPLY---RYIQWLRLRRASWRLAFNPQDKVIDIALDAGFQNPE 93
AlbA Klebsiella       SWVQTLELMKMYDRWFSQQELQVLPFAEQDEQRNQTWLELVGEAQQLM------GERCP- 173
                      :  . :.: : :  * .        : :  . *. :*   ..  ::::      * : *

STM3175 Salmonella    SFTRAFKTAFGQSPRRFRQSPDWLAWHQRV----PKL--ALQEQHVMDVKIVE---FPPT 144
AlbA Klebsiella       ----------ADEPRAIALATRWMEQLEQDTAGRPEFLTRLNEMHAAEPQMREQTGVTPE 223
                                .:.** :  :  *:   ::     *::   *:* *. : :: *   . *

STM3175 Salmonella     RVAMLTHLGHPDKVNASAAKFIAWRRETGQSPIASSQ--TFG--IAWHDPQTTPPAQFRF 200
AlbA Klebsiella        MIDFITR-------AFAESKLAIWARYLNAEELAFTRQHYFDRLMEWPALVA-----DLH 271
                        : ::*:         : :*:  * *  . . :* ::   *.  : *    :       .

STM3175 Salmonella     DICGSVRQPIAENDVGVVNSEIPGGRCAVVRHQGSLDSLPESVWY-LFREWLPASGETPR 259
AlbA Klebsiella        RACREKRDPASP-----------EGQQ-----------LA-QRWLALFQSYAGKDAQTQQ 308
                         * . *:* :             *:            *  . *  **:.:   ..:* :

STM3175 Salmonella     DFPVFFQYLNFVHEVAEHELLTDIYLPLR--------------------- 288
AlbA Klebsiella        KFR----Y---AMEQ-EPHLMKGTWMTSEVLSWLQQAIGVMMRQAQGPAAE 351
                       .*     *   . *  * .*:.. ::  .
```

**Fig. S8**. **Domain structure and sequences of STM3175 and AlbA.** A) Domains identified by the NCBI conserved domains search tool (*2*) are highlighted in bold. B) CLUSTAL Omega sequence alignment of STM3175 and AlbA.

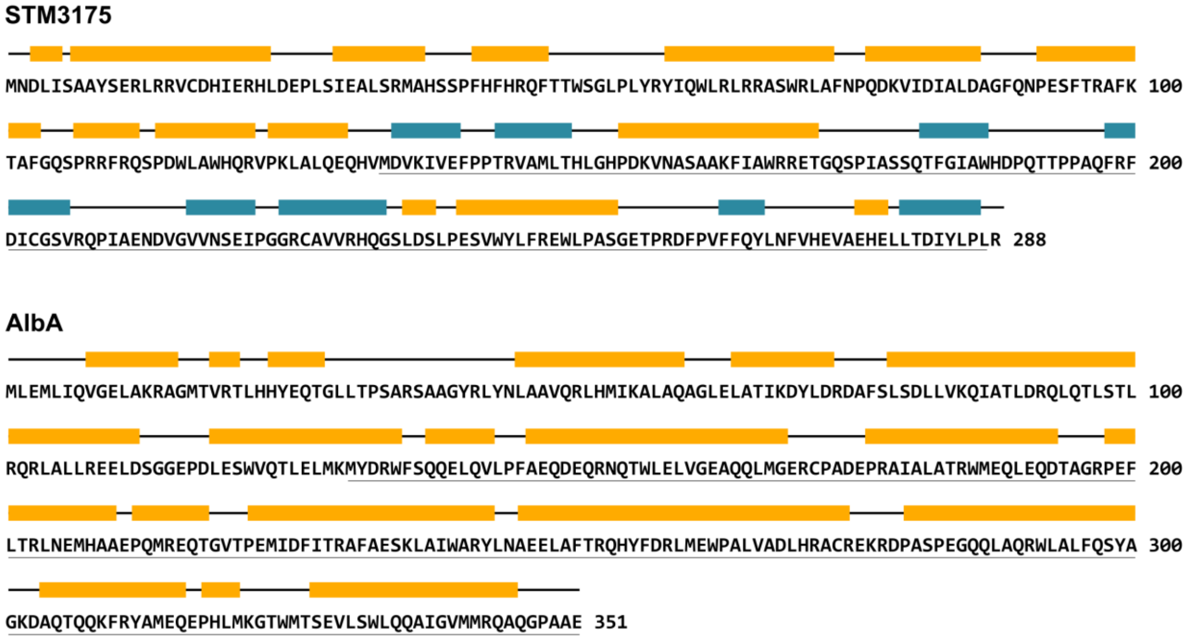

**Fig. S9**. **Secondary structures of STM3175 and AlbA.** Secondary structures predicted from the amino acid sequences by the PSIPRED Server. α-helices are colored in yellow, β-sheets are shown in blue. Amino acid residues in the ligand-binding domains are underlined.

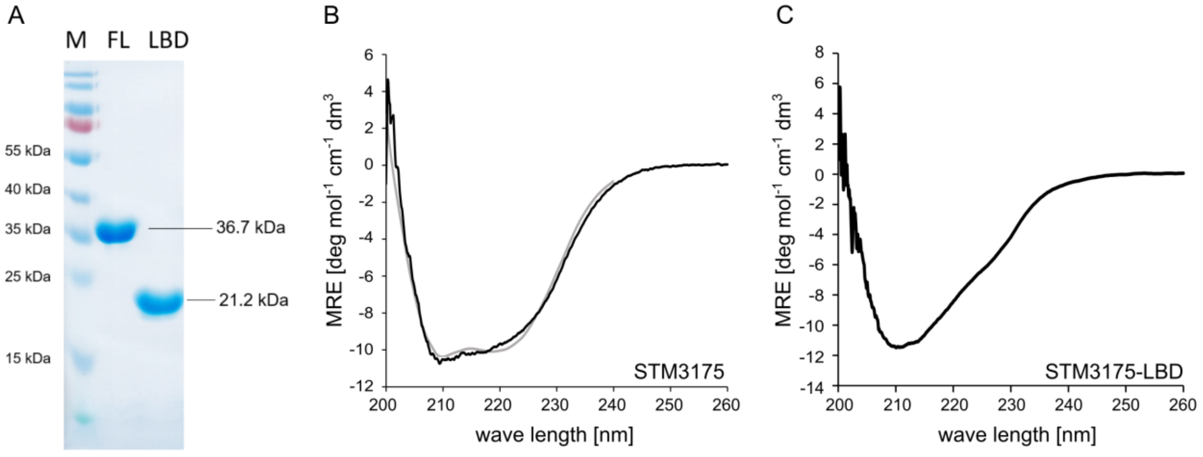

**Fig. S10**. **Circular dichroism spectroscopy of STM3175**. A) SDS-PAGE with $His_6$-STM3175 (FL) and $His_6$-STM3175-LBD (LBD) after Ni-NTA purification. B) CD spectrum of $His_6$-STM3175 in Tris buffer (black) and with K2D reconstructed spectrum (grey). C) CD spectrum of the ligand binding domain in Tris buffer. MRE = mean residue ellipticity.

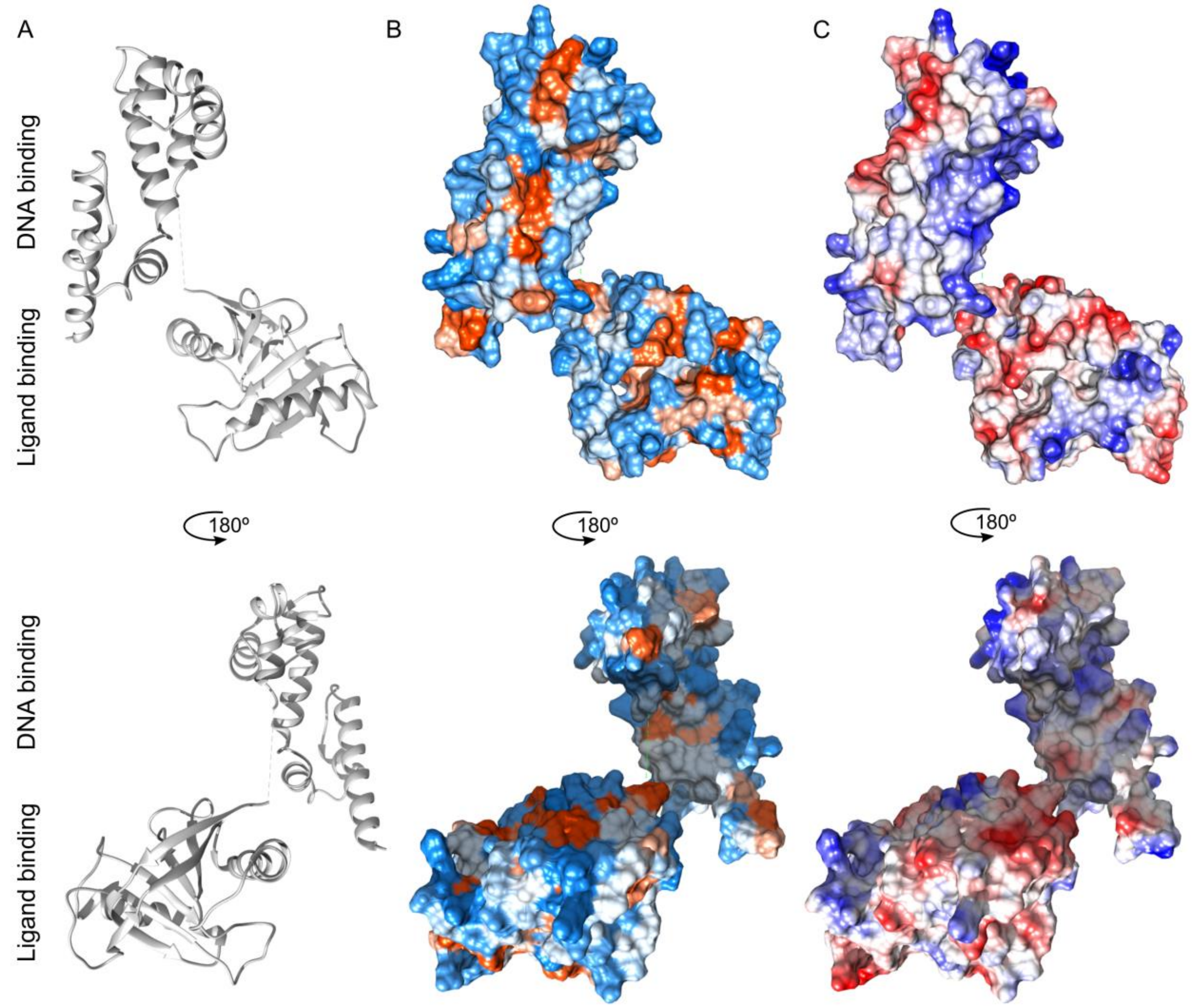

**Fig. S11. STM3175 surface analysis.** A) STM3175 crystal structure with B) hydrophobic residues colored in orange and polar amino acids shown in light blue and C) negatively charged amino acids in red and positively charges residues in blue.

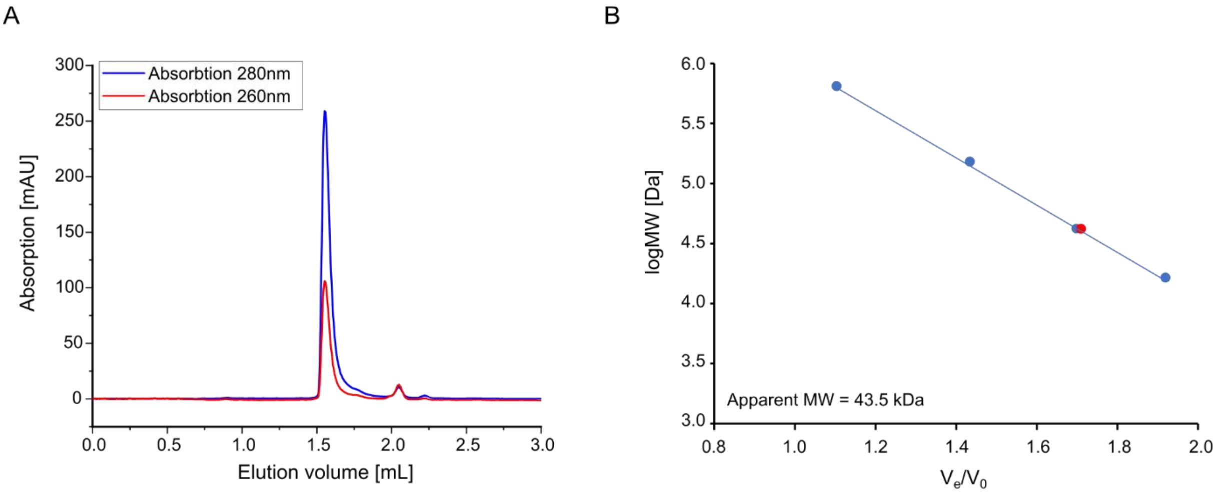

**Fig. S12 Analytical size exclusion chromatography**. A) Chromatogram of an analytical size exclusion gel filtration run of STM3175 on a Superdex S200 increase 3.2/300. STM3175 (with a concentration of 30 µM in 50 µl in gel filtration buffer) was analysed with a flow rate of 0.04 ml/min. B) Calibration curve of the column with thyroglobulin (670 kDa), γ-globulin (158 kDa), ovoalbumin (44 kDa) and myoglobin (17 kDa) shown as blue circles. The apparent molecular weight of STM3175 was calculated from the elution volume using the linear regression (blue line). The apparent molecular weight of STM3175 is 43.5 kDa (data point shown in red).

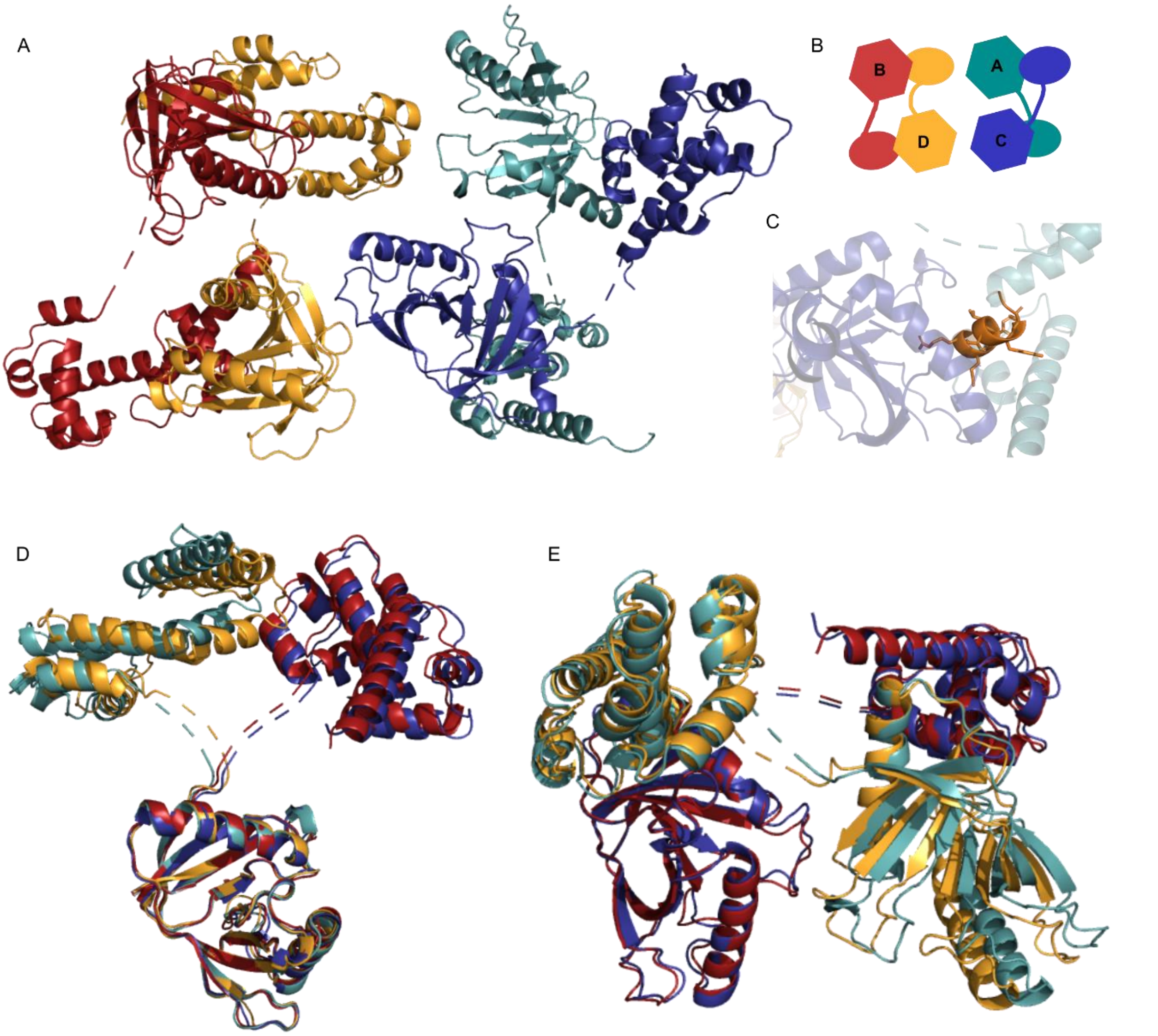

**Fig. S13. Spatial organization of STM3175 in the crystal structure.** A) Arrangement of two domain-swapped dimers in the crystal. B) schematic of the monomer organization in a unit cell with hexagons symbolizing the ligand binding domains and ellipses as HTH domains. C) A helix (highlighted in orange) of the DBD of monomer A occupies the C-terminal end of the binding groove on the LBD of monomer C. D) Alignments of the LBD of the four monomers show two distinct domain orientations. Monomers A and D resemble each other and monomers

B and C align. E) each dimer consists of two monomers with different conformations, but alignments show that the two dimers are not identical.

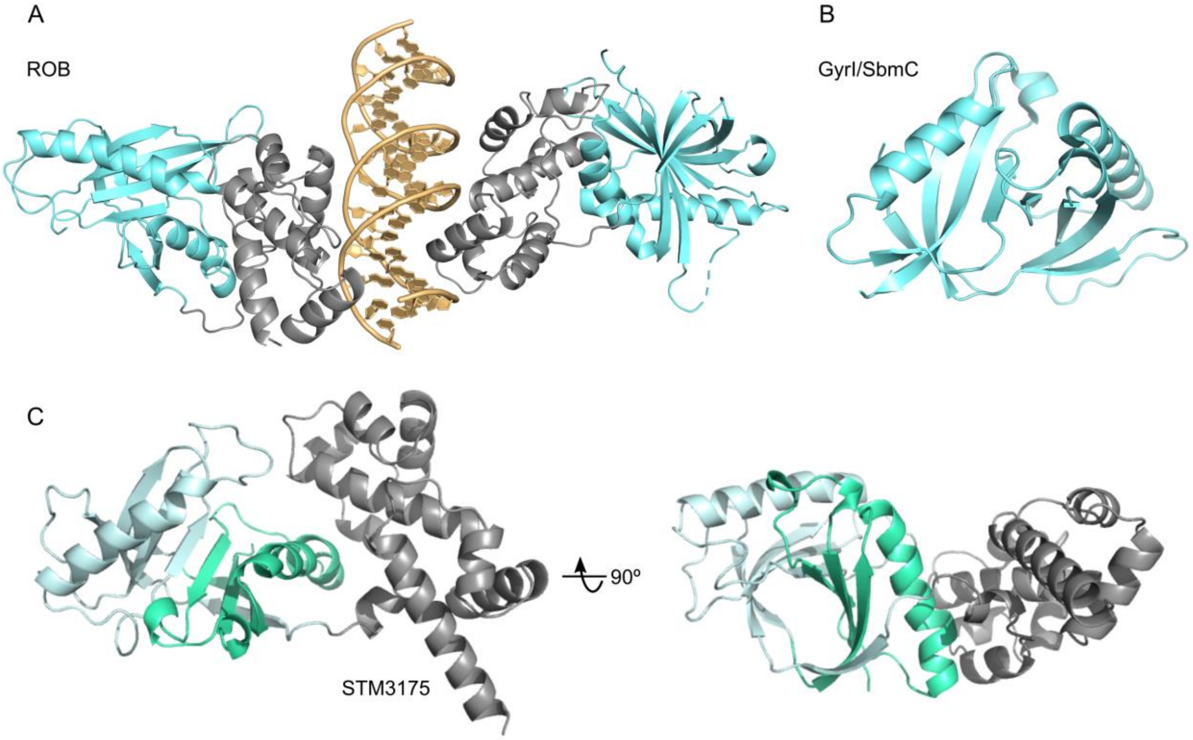

**Fig. S14**. **STM3175 homology model and modeling templates.** A) Crystal structure of the highest scoring template, transcription factor ROB, bound to its cognate DNA in a tertiary complex (PDB-ID: 1d5y). B) Crystal structure of the highest scoring template for the ligand-binding domain SbmC (GyrI; PDB-ID: 1jyh). C) Robetta homology model of STM3175. The DBDs are colored in grey, LBD in cyan. In STM3175, SH2 subunits of the pseudo-dimeric motif are colored in pale cyan and green.

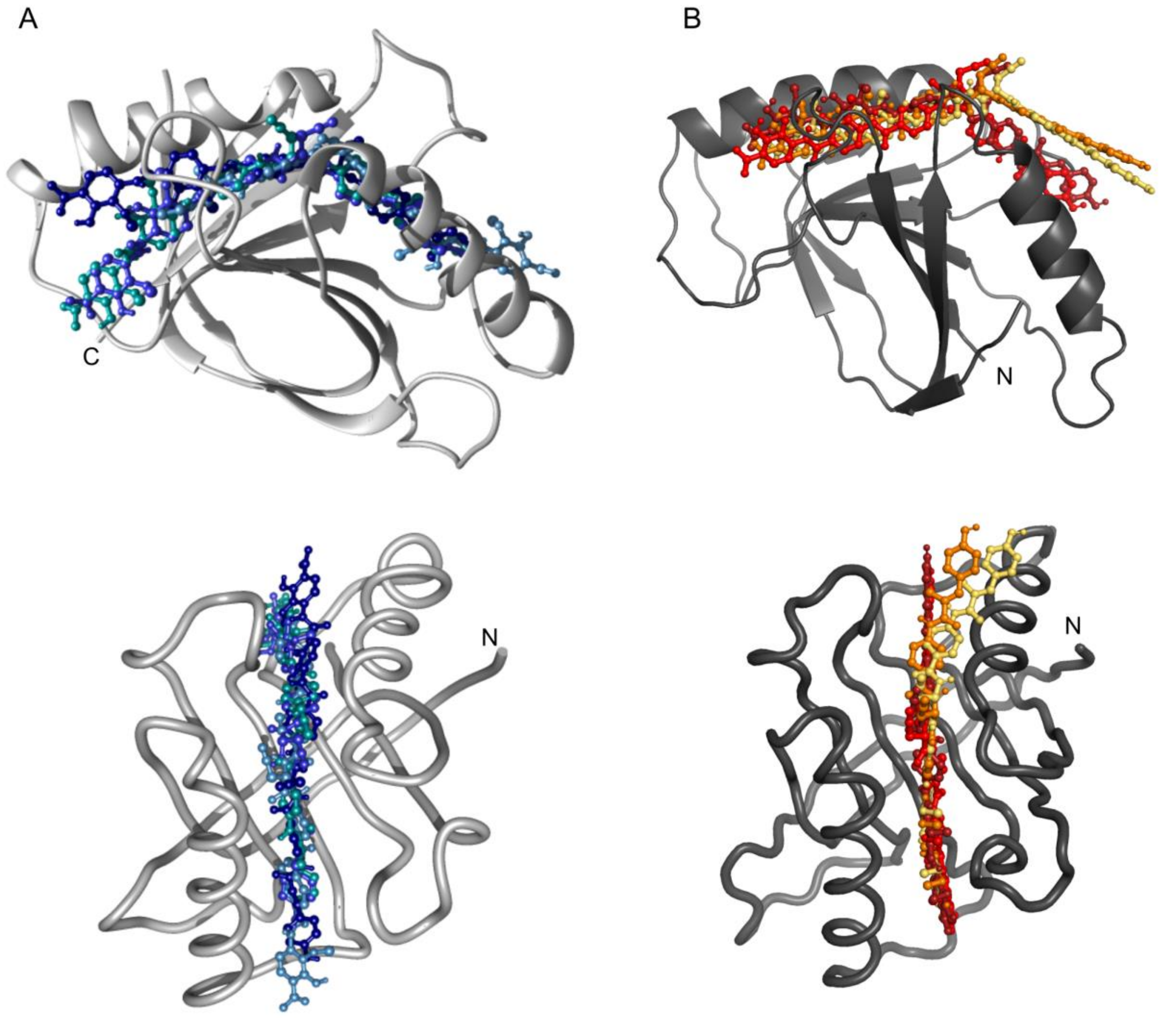

**Fig. S15.** S LBDs of STM3175 and *E. coli* YgiV with albicidin. A) Conformation of albicidin in the four best-ranked models in AutoDock Vina with the crystal structure of STM3175. The top-ranked conformation is shown in dark blue. B) Rosetta model of *E. coli* YgiV docked with albicidin using HADDOCK. The four best-ranked models are shown with the top-ranking albicidin conformation shown in dark red.

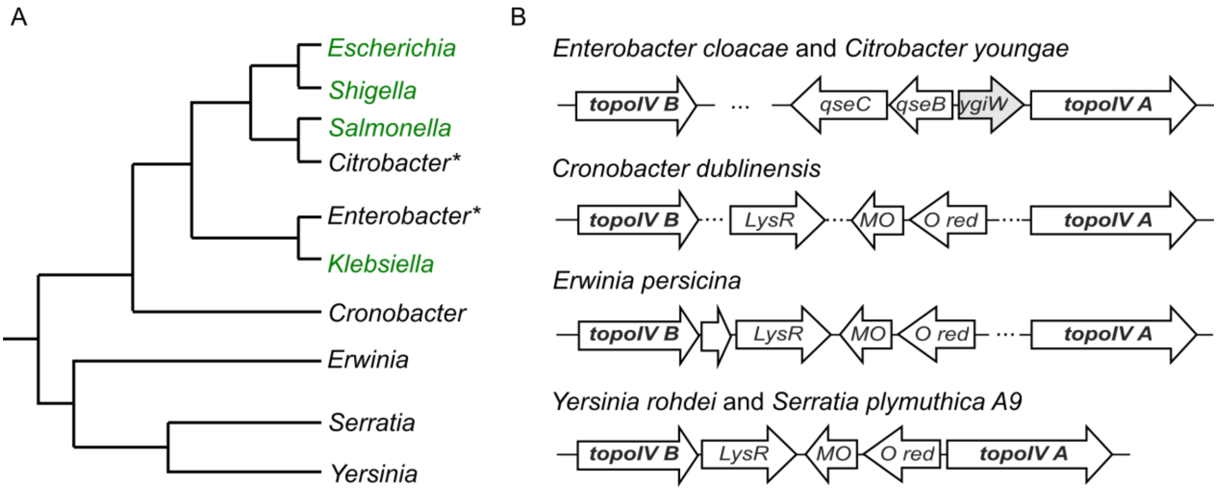

**Fig. S16. Enterobacteriaceae family tree and genomic context of topoisomerase IV subunits.** A) Genera with mono- or di-domain STM3175 homologs in the same genomic context are highlighted in green. Genera where YgiW was present but no STM3175 homolog are indicated with an asterisk. The phylogenetic tree was adapted from Hata et al. (*10*). B) Genomic context of the two subunits of topoisomerase I, *topoIV A* and *topoIV B*. *ygiW* is shown in grey. *LysR* = LysR family transcriptional regulator; *MO* = antibiotic biosynthesis monooxygenase; *O red* = NAD(P)H-dependent oxidoreductase. Genomic Sequences were obtained from the NCBI database with the following accession numbers: *E. cloacae:* NZ_CP009756.1, *C. youngae*: NZ_GG730303.1, *Y. rohdei*: NZ_CP009787, *S. plymuthica A9*: NC_015567.1, *C. dublensis*: NZ_CP012266.1, *E. persicina*: NZ_CP082141.1.

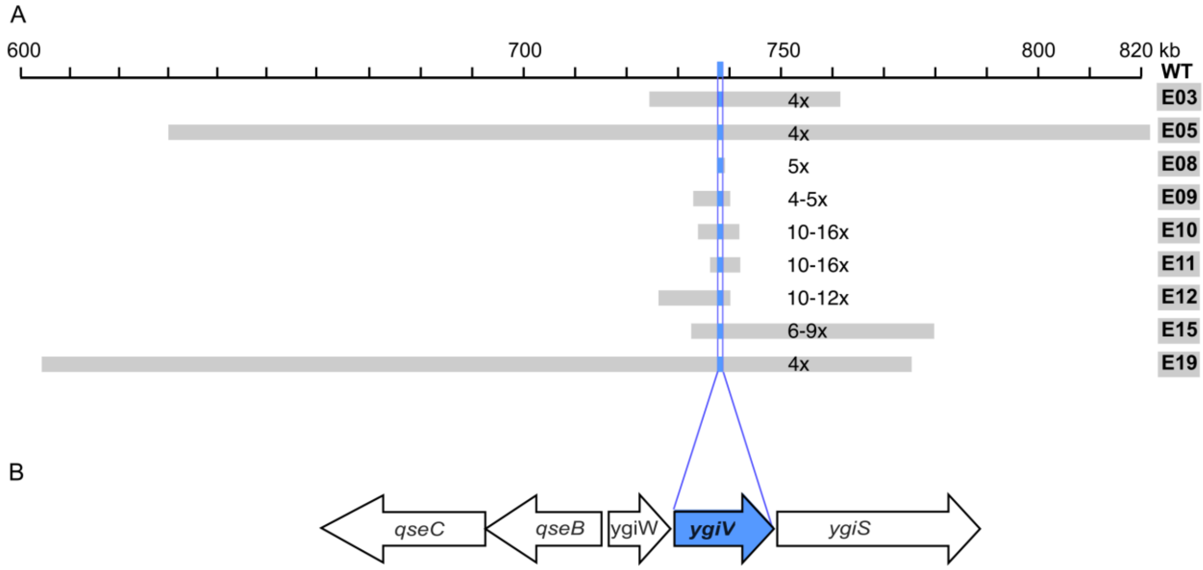

**Fig. S17. Gene duplication and amplification (GDA) in *E. coli* strains with *a*lbicidin hyper-resistance. A)** Mapping of GDA-strains after treatment with albicidin (increasing albicidin conc. from 01.0156 µg $mL^{-1}$ to 8 µg mL$^{-1}$ in ten passages), which are different in copy number and size (Table S2). B) The common ~600 bp long region (blue) includes the gene *ygiV*.

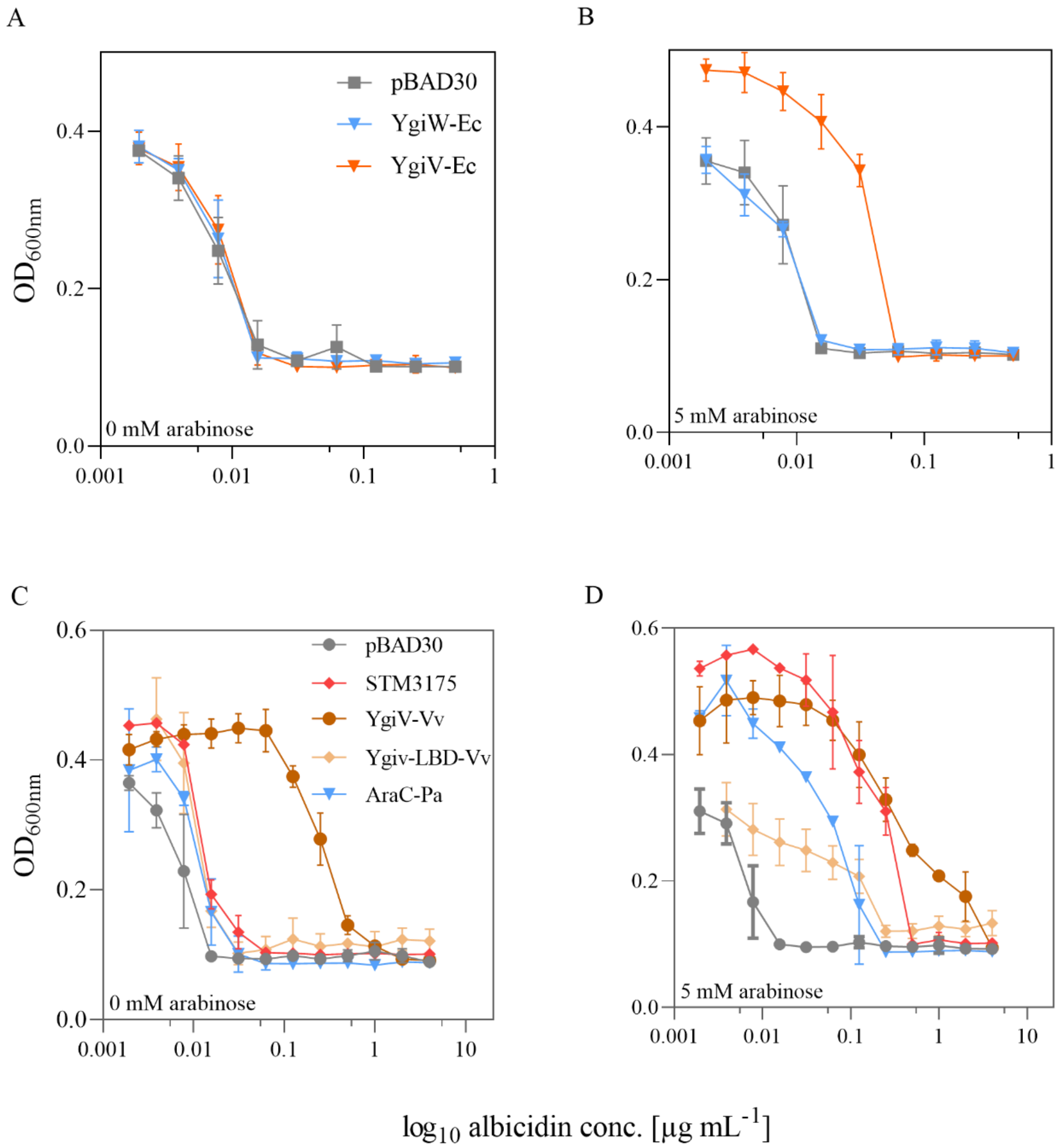

**Fig. S18. Comparison of non-induced and arabinose induced expression systems of STM3175 homologs from *E. coli*, *V. vulnificus* and *P. aeruginosa* in *S.* Typhimurium.** A-B) Arabinose induction results in elevated albicidin MIC in *S*. Typhimurium cells where the homolog YgiV-Ec from *E. coli* is expressed but not when YgiW-Ec is expressed. C-D) Arabinose induced over-expression of the STM3175 homologs YgiV-Vv from *Vibrio vulnificus,* its LBD (YgiV-LBD-Vv) and AraC-Pa from *Pseudomonas aeruginosa* leads to increased albicidin MICs. The increased MIC for YgiV-Vv even without arabinose induction is due to its DBD, which is an AraC-homolog and might results in autoregulation under albicidin treatment. Sensitivity of Ygiv-Vv to albicidin without arabinose induction is restored after cloning of only LBD of YgiV. The data represent means and standard deviations of three biological replicates that were performed in three technical replicates, except YgiW-Ec, which was performed in two biological and three technical replicates.

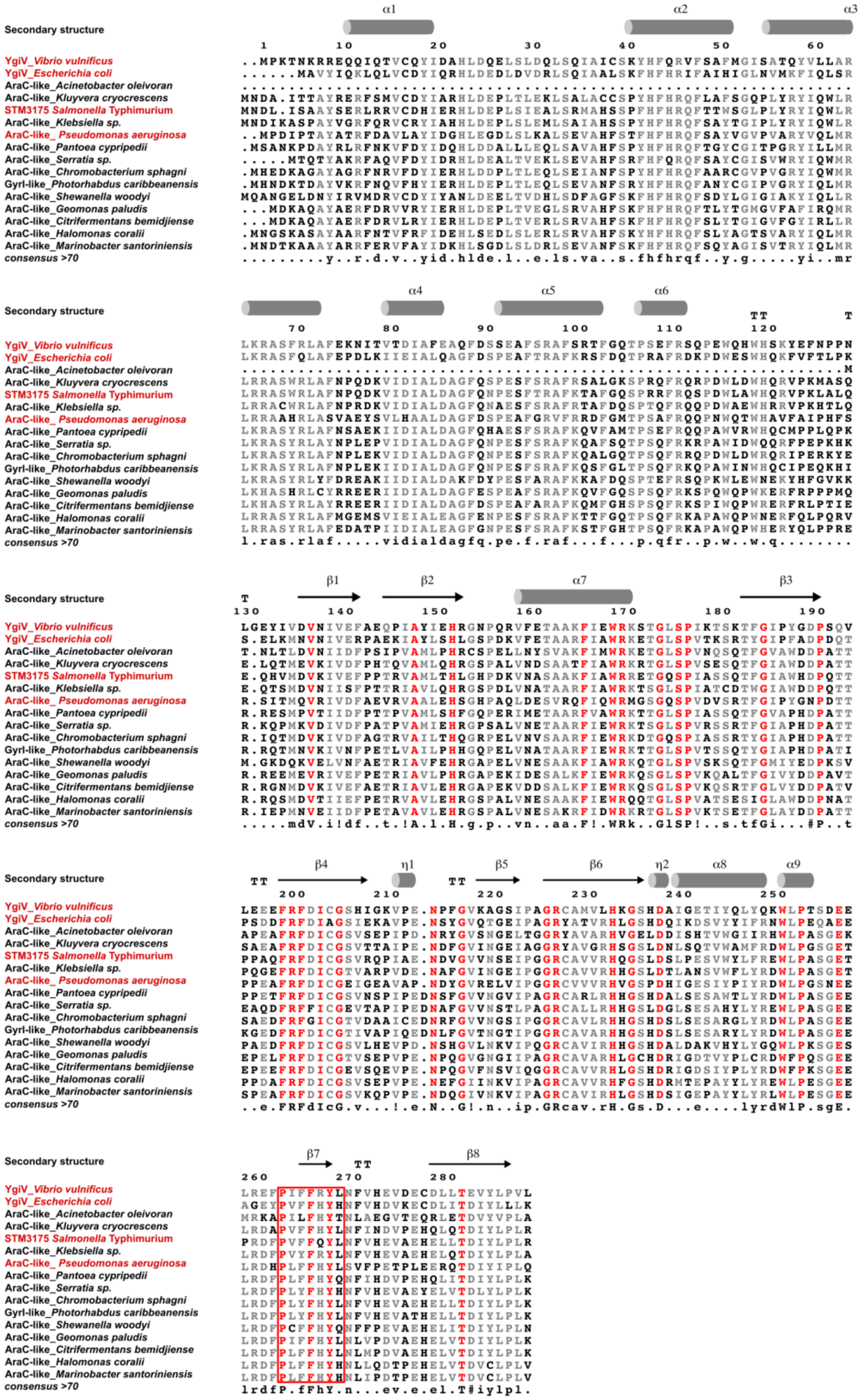

**Fig. S19. Alignments of STM3175 homologs.** Shown are representatives from different genera of the top 100 homologs identified by BLAST of the LBD against the RefSeq database excluding *Salmonella* sp.. Proteins investigated in this study are indicated in red. Amino acids with 100% consensus are shown in red and the beta-strand forming the base of the binding groove is highlighted by a red box.

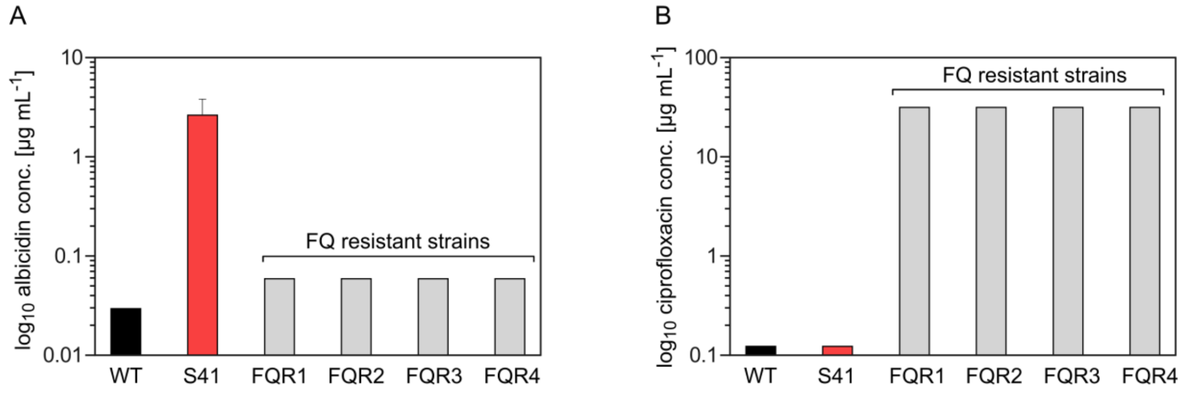

**Fig. S20. MICs of albicidin and ciprofloxacin**. Determination of the MIC for the wild type (WT, ATCC 14028), the albicidin evolved strain (S41) and 4 high fluoroquinolones (FQ) resistant strains (FQR1-4) for A) albicidin and B) ciprofloxacin in 96-well-plates. The error bars represent mean and standard deviations three biological replicates that were performed in three technical replicates (except FQR1-4 for ciprofloxacin, which represent mean and standard deviations three technical replicates). Strains without error bar have the same result in every single experiment.

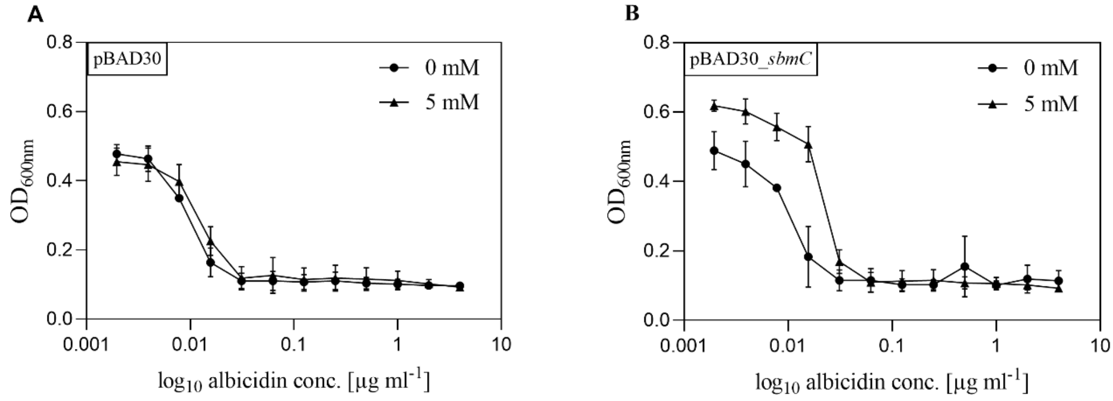

**Fig. S21. Comparison of arabinose induced expression system of *S.* Typhimurium SbmC and empty vector in *S.* Typhimurium**. A) MIC determination without arabinose and 5 mM arabinose in empty vector control pBAD30, B) in pBAD30::*sbmC*. The error bars represent mean and standard deviations of 2 biological replicates that were performed in 3 technical replicates.

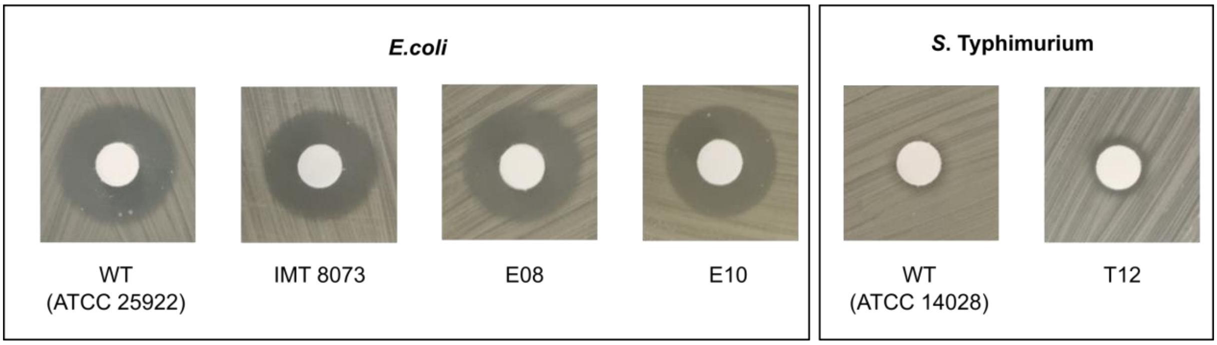

**Fig. S22. Agar diffusion assay with MccB17 and evolved *E. coli* and *S.* Typhimurium strains.** At concentrations of 10 mg/ml, no difference in susceptibility towards MccB17 was observed between WT or evolved *E. coli*

strains E08 and E10. *S*. Typhimurium WT strain was not susceptible to MccB17 and neither was the evolved strain T12.

Table **S1. Genetic changes in albicidin resistant *Salmonella* isolates.**

| Strain | SNPs/ deletions |
|---|---|
| S08 | Tsx, Trp146*; AegA, Val438Gly |
| S09 | Tsx, Trp146* |
| S10 | Tsx, 380del |
| S11 | Tsx, Trp242* |
| S12 | Tsx, Thr19fs |
| S16 | Tsx, Thr225fs |
| S27 | Tsx, Asp221fs |
| S36 | Tsx, Asn223fs |
| S39 | Tsx, Arg187fs |
| S45 | Tsx, Gln37fs |
| S18 | SSU ribosomal protein S1, Tyr205Cys |
| S19 | QseC, Arg12fs |
| S20 | - |
| S41 | - |
| S47 | CCA tRNA nucleotidyl transferase, Leu322Arg |
| Abbreviations: | *, stop codon; del, deletion; fs, frameshift |

**Table S2.** Summary of evolved *S.* Typhimurium and *E. coli* strains harboring a GDA-region

| | GDA – region | | | Albicidin |
|---|---|---|---|---|
| Strain ID<br>*S.* Typhimurium | Genomic region | Copy number | Size [kB] | Max. detected conc. [µg $mL^{-1}$] |
| **S41 (**8640-41) | 537.415 – 584.697 | 3 – 4 | 47 | 2 |
| **T01 (**9866-01) | 500.000 – 658.112 | 5 | 158 | 20 |
| **T04 (**9866-04) | 456.463 – 546.76 | 3 – 4 | 90 | 20 |
| **T05** (9866-05) | 523.665 – 665.000 | 4 | 141 | 20 |
| **T10 (**9866-10) | 543.867 – 546.804 | 10 – 15 | 3 | 20 |
| **T12 (**9866-12) | 544.528 – 550.256 | 7 | 6 | 20 |
| Strain ID<br>***E. coli*** | Genomic region | Copy number | Size [kB] | Max. detected conc. [µg $mL^{-1}$] |
| **E19** (25922-19) | 605.320 – 775.048 | 4 | 170 | 8 |
| **E05** (25922-05) | 630.307 – 822.228 | 4 | 192 | 8 |
| **E08** (25922-08) | 738.859 – 739.503 | 5 | 0.6 | 8 |
| **E03** (25922-03) | 725.441 – 762.165 | 4 | 36.7 | 8 |
| **E15** (25922-15) | 732.124 – 779.032 | 6 – 9 | 47 | 8 |
| **E10** (25922-10) | 735.358 – 742.724 | 10 – 16 | 7.4 | 8 |
| **E09** (25922-09) | 734.500 – 740.630 | 4 – 5 | 6.1 | 8 |
| **E11** (25922-11) | 737.160 – 742.810 | 10 – 16 | 5.6 | 8 |
| **E12** (25922-12) | 726.298 – 740.552 | 10 - 12 | 1.4 | 8 |

Table **S3. Genetic changes in albicidin resistant *Δtsx Salmonella* isolates.**

| Strain | GDA | SNPs/ deletions | MIC [µg $mL^{-1}$] |
|---|---|---|---|
| T01 | + | **-** | ≥ 20 |
| T04 | + | QseB, Ile49Met | ≥ 20 |
| T05 | + | STM3175, Ile5Asn | ≥ 20 |
| T10 | + | - | ≥ 20 |

| | | | |
|---|---|---|---|
| T12 | + | Topoisomerase IV subunit B, Asp476Glu; FIG01045621 hypothetical protein, Thr30Ala, 110kb deletion fragment | ≥ 20 |

**Table S4.** Summary of evolved *S.* Typhimurium strain T12 after growth without antibiotic pressure

| | | GDA - region | | Albicidin |
|---|---|---|---|---|
| Strain ID *S.* Typhimurium | Subcultivation | Copy number | Size [kB] | 1xMIC [µg $mL^{-1]}$ |
| **T12** (input) | 0 | 7 | 6 | ≥20 |
| **T12-1** | 5 | 7 | 6 | ≥8 |
| | 6 | 2 | 6 | 4 |
| | 15 | 2 | 6 | 4 |
| **T12-2** | 15 | 7 | 6 | ≥8 |
| **T12-3** | 15 | n.d | n.d | ≥8 |

**Table S5.** MIC values of STM3175-LBD of non-induced and induced arabinose expression in different strains

| arabinose | 0 mM | 5 mM |
|---|---|---|
| **WT** | µg $mL^{-1}$ | |
| pBAD30 | 0,0156 | 0,0156 |
| STM3175-LBD | 0,03 | 0,06 |
| **WT Δ*STM3175*** | | |
| STM3175-LBD | 0,0156 | 0,06 |

**TableS6.** Antibiotic susceptibility testing of wild type strain (WT) and the respective evolved strain (S41) for 24 different antibiotics.

| **Strains** | **WT** | **S41** |
|---|---|---|
| **Antibiotic** | 1 x MIC [µg $mL^{-1}$] | |
| Amoxicillin/ clavulanic acid (2:1 ratio) | 1/0.5 | 1/0.5 |
| Ampicillin | 1 | 1 |
| Cefoperazone | 0.5 | 0.5 |
| Cefotaxime | 0.06 | 0.12 |
| Cefquinome | 0.06 | 0.06 |
| Ceftiofur | 0.5 | 1 |
| Cephalothin | 2 | 2 |
| Ciprofloxacin | 0.12 | 0.12 |
| Colistin | R (> 256) | R (> 256) |
| Doxycycline | 0.5 | 0.5 |
| Enrofloxacin | 0.5 | 0.25 |

| | | |
|---|---|---|
| Florfenicol | 0.25 | 0.25 |
| Gentamicin | 0.5 | 0.5 |
| Imipenem | >128 (R) | >128 (R) |
| Marbofloxacin | 0.25 | 0.25 |
| Nalidixic Acid | 16 | 8 |
| Neomycin | 4 | 4 |
| Penicillin | 16 | 16 |
| Streptomycin | 2 | 2 |
| Tetracycline | 1 | 1 |
| Tiamulin | 1 | 1 |
| Tilmicosin | 128 | 128 |
| Trimethoprim/ sulfamethoxazole | 0.06/1.19 | 0.06/1.19 |
| Tulathromycin | 8 | 8 |

Abbreviations: **R, resistant**

**TableS7.** Antibiotic susceptibility testing of Δ*tsx* mutant strain and the respective evolved strains for 24 different antibiotics.

| **Strains** | **Δtsx mutant** | **T01** | **T04** | **T05** | **T10** | **T12** |
|---|---|---|---|---|---|---|
| **Antibiotic** | 1 x MIC [μg $mL^{-1}$] | | | | | |
| Amoxicillin/ clavulanic acid 2:1 ratio | 1/0.5 | 1/0.5 | 1/0.5 | 1/0.5 | 1/0.5 | 1/0.5 |
| Ampicillin | 1 | 1 | 1 | 1 | 1 | 1 |
| Cefoperazone | 0.5 | 0.5 | 0.5 | 0.5 | 0.5 | 0.25 |
| Cefotaxime | 0.06 | 0.12 | 0.06 | 0.12 | 0.12 | 0.06 |
| Cefquinome | 0.06 | 0.06 | 0.12 | 0.06 | 0.12 | 0.06 |
| Ceftiofur | 0.5 | 1 | 1 | 1 | 1 | 1 |
| Cephalothin | 2 | 2 | 2 | 2 | 2 | 2 |
| Ciprofloxacin | 0.25 | 0.25 | 0.25 | 0.25 | 0.25 | 0.25 |
| Colistin | 0.5 | 0.5 | 1 | 1 | 4 | 1 |
| Doxycycline | 2 | 2 | 2 | 2 | 1 | 2 |
| Enrofloxacin | 0.5 | 0.5 | 0.5 | 0.5 | 0.5 | 0.5 |
| Florfenicol | 4 | 8 | 8 | 8 | 4 | 4 |
| Gentamicin | 0.5 | 0.25 | 0.5 | 0.5 | 0.5 | 0.25 |
| Imipenem | 0.25 | 0.25 | 0.25 | 0.25 | 0.12 | 0.25 |
| Marbofloxacin | 0.25 | 0.25 | 0.25 | 0.25 | 0.25 | 0.25 |
| Nalidixic Acid | >256 (R) | >256 (R) | >256 (R) | >256 (R) | >256 (R) | >256 (R) |
| Neomycin | 0.5 | 0.5 | 0.5 | 0.5 | 0.5 | 0.5 |
| Penicillin | 16 | 16 | 8 | 16 | 16 | 8 |
| Streptomycin | 16 | 16 | 16 | 8 | 8 | 8 |
| Tetracycline | 2 | 2 | 2 | 2 | 1 | 2 |
| Tiamulin | >128 (R) | >128 (R) | >128 (R) | >128 (R) | >128 (R) | >128 (R) |
| Tilmicosin | 128 | >256 | >256 | >256 | 128 | 128 |
| Trimethoprim/ sulfamethoxazole | 0.06/1.19 | 0.06/1.19 | 0.06/1.19 | 0.06/1.19 | 0.06/1.19 | 0.03/0.59 |
| Tulathromycin | 8 | 8 | 16 | 16 | 8 | 8 |

Abbreviations: **R, resistant**

**TableS8.** Susceptibility testing against microcin B17 of *E. coli* and *S.* Typhimurium strains

| Strain | 1xMIC microcin B17 [μg $mL^{-1}$] | Agar diffusion microcin B17 [mm] |
|---|---|---|
| *E.coli* IMT 8073* | 4 | 18 |
| *E.coli* ATCC 25922 | 4 | 18 |
| *E.coli* E08 | 4 | 18 |
| *E.coli* E10 | 4 | 18 |
| *S.* Typhimurium WT | ≥ 128 | 8 |
| *S.* Typhimurium T12 | ≥ 128 | 8 |

*contains *SbmC* gene

**TableS9.** Strains and plasmids used in this work

| Strain | Relevant Features | Reference/ Source |
|---|---|---|
| 8640 | *S.* Typhimurium ATCC 14028 wild type, virulent, NalR | laboratory stock |
| 9866 | *S.* Typhimurium ATCC 14028 NalR Δtsx | this study |
| 11078 | *S.* Typhimurium ATCC 14028 NalR ΔqseBC | this study |
| 11534 | *S.* Typhimurium ATCC 14028 NalR ΔygiW-STM3175 | this study |
| 11536 | *S.* Typhimurium ATCC 14028 NalR Δtsx ΔygiW-STM3175 | this study |
| 11656 | *S.* Typhimurium ATCC 14028 NalR Δtsx ΔqseBC | this study |
| 2100 | *S.* Typhimurium LT2 strain JR501 hsdSA29 hsdSB121 hsdL6 (r- m+) | SGSC |
| 1948 | *E. coli* DH5 | laboratory stock |
| MG1655 | *E. coli* MG1655 | laboratory stock |
| 25922 | *E. coli* ATCC 25922 wild type | laboratory stock |
| BL21 | *E. coli* BL21-Gold (λDE3) | AG Süßmuth, TU Berlin |
| IMT8073 | *E. coli* | laboratory stock |
| FQR1 | *S.* Typhimurium IMT42052-6 $bla_{TEM,}$ catA1, aph(3′)-Ia, ant(3″)-Ia, sul1, tet(B), dfrA5/A14, dfrA1/A15/A16 | (*24*) |
| FQR2 | *S.* Typhimurium IMT42053-7 $bla_{TEM,}$ catA1, aph(3′)-Ia, ant(3″)-Ia, sul1, tet(B), dfrA5/A14, dfrA1/A15/A16 | (*24*) |
| FQR3 | *S.* Typhimurium IMT42054-8 $bla_{TEM,}$ catA1, aph(3′)-Ia, ant(3″)-Ia, sul1, tet(B), dfrA5/A14, dfrA1/A15/A16 | (*24*) |
| FQR4 | *S.* Typhimurium IMT42055-15 $bla_{TEM,}$ catA1, aph(3′)-Ia, ant(3″)-Ia, sul1, tet(B), dfrA5/A14, dfrA1/A15/A16 | (*24*) |

| Plasmids | Relevant Features | Reference/ Source |
|---|---|---|
| pBAD30 | bla araC+ ParaBAD p15Aori | Guzman et al., 1995(*25*) |
| pBAD30_qseB | bla araC+ ParaBAD–qseB+ p15Aori | this study |
| pBAD30_ygiW | bla araC+ ParaBAD –ygiW+ p15Aori | this study |
| pBAD30_STM3175 | bla araC+ ParaBAD–STM3175+ p15Aori | this study |
| pBAD30_STM3175-DB | bla araC+ ParaBAD–STM3175+ΔR110-R288 p15Aori | this study |
| pBAD30_STM3175-LBD | bla araC+ ParaBAD–STM3175+ ΔM0-R110 p15Aori | this study |
| pBAD30_ygiV | bla araC+ ParaBAD–ygiV+ p15Aori | this study |
| pBAD30_ygiW | bla araC+ ParaBAD–ygiW+ p15Aori | this study |
| pBAD30-ygiV-Ec | bla araC+ ParaBAD–ygiV-Ec+ p15Aori | this study |
| pBAD30-ygiW-Ec | bla araC+ ParaBAD–ygiW-Ec+ p15Aori | this study |
| pET28a(+) | KanR lacI+ PT7-lacO pBR322ori | Novagen |
| pET28a(+)_STM3175 | KanR lacI+ PT7-lacO-STM3175+ pBR322ori | this study |
| pET28a(+)_STM3175-DB-P114 | KanR lacI+ PT7-lacO-STM3175+ ΔP114-R288 pBR322ori | this study |
| pET28a(+)_STM3175-DB-V133 | KanR lacI+ PT7-lacO-STM3175+ ΔV133-R288 pBR322ori | this study |
| pET28a(+)_STM3175-LBD-P114 | KanR lacI+ PT7-lacO-STM3175+ ΔM0-P114 pBR322ori | this study |
| pET28a(+)_STM3175-LBD-V133 | KanR lacI+ PT7-lacO-STM3175+ ΔM0-V133 pBR322ori | this study |
| pET28a(+)-ygiV | KanR lacI+ PT7-lacO-ygiV+ pBR322ori | this study |
| pUC57-ygiV-Vv | bla($Ap^R$) lacZ ygiV-Vv+ pMB1ori | GenScript |
| pUC57-araC-Pa | bla($Ap^R$) lacZ araC-Pa+ pMB1ori | GenScript |
| pBAD30-ygiV-Vv | bla araC+ ParaBAD–ygiV-Vv+ p15Aori | this study |
| pBAD30-ygiv-LBD-Vv | bla araC+ ParaBAD–ygiV-LBD-Vv+ p15Aori | this study |
| pBAD30-araC-Pa | bla araC+ ParaBAD–araC-Pa+ p15Aori | this study |

Abbreviations: $Nal^R$, nalidixic acid-resistance; $Cb^R$ , carbenecellin-resistance; $Kan^R$, kanamycin-resistance; $Ap^R$, ampicillin-resistance; SGSC, Salmonella Genetic Stock Centre; DB, DNA binding domain; LBD, ligand-binding domain; Ec, *Escherichia coli;* Vv, *Vibrio vulnificus*; Pa, *Pseudomonas aeruginosa*

**TableS10.** Primer with their sequence and target region used in this work

| Primer | Sequence | Target region |
|---|---|---|
| YGIWXbalF | GATCATGCTCTAGATGAAAGGGAAAAGTAATCATGAAAAAATT-AGCTG | ygiW ORF |
| YGIWHind3R | GATCATGCAAGCTTGACCGATCTTGCGCAATGTGGGATTACGGAT-TCAC | ygiW ORF |
| 3175XbalF | GATCATGCTCTAGAGGCCACAAGGAGGCAGTGATGAATGAC | STM3175 ORF |
| 3175Hind3R | ATGCGATTGGTCTGAGTCACAAAGCTTGAGACAGG | STM3175 ORF |
| PREAEcoRIF | GGCAACGCGAATTCCCGCAAGGAAGAACAGATGCGA | qseB gene |
| PREAXbalR | AGCTCAGGTCTAGACGTTGCGTCAATTTCATGCGTCAC | qseB gene |
| 3175-Xho-AraC-F2 | AGTCGTACCTCGAGCTTCCTCGCCACCTGTC | STM3175-DB gene |
| 3175-Xho-AraC-R | ATGGCCGATCTCGAGAAACCGGCGCGGG | STM3175-DB gene |
| 3175XhoIF | AGTCGTACCTCGAGTTTCGGCAATCGCCG | STM3175-LBD gene |
| 3175XhoIR | ATGGCCGATCTCGAGCATCACTGCCTCC | STM3175-LBD gene |
| 3175NdeIF | AGTCGTACCATATGAGCGGCGAAAACCTG-TATTTTCAGGGCGCTAGCATGAATGAC-CTGATCAGCGCGGCTTATTCCG* | STM3175 gene |
| 3175BamHIR | ATGCCGATGGATCCTGAGACAGGTGGCGAGGAAG | STM3175 gene |
| 3175NdeI_VF | AGTCGTACCATATGAGCGGCGAAAACCTG-TATTTTCAGGGCGCTAGCATGGTCATGGACGTAAAAATCGTTG* | STM3175 gene |
| 3175BamHI_VR | GCATGCATGGATCCTTAGACATGCTGCTCCTGTAACG | STM3175 gene |
| 3175NdeI_PF | AGTCGTACCATATGAGCGGCGAAAACCTG-TATTTTCAGGGCGCTAGCATGCCG-GACTGGCTGGCCTGGCACCAGCGC* | STM3175 gene |
| 3175BamHI_PR | GCATGCATGGATCCTTACGGCGATTGCCGAAACCGGCGC | STM3175 gene |
| SBMCXbaIF | GATCGATCTCTAGACATTGTGAAGTGACGGAGGCAGCATG | sbmC gene |
| SBMCHind3R | GCATGCATAAGCTTAGGAAACCAGTCGATCACTTCCG | sbmC gene |
| TsxF | ATAGGCTCCGCAGAAACACG | tsx gene |
| TsxR | GCCGGAAGTAATGTGAAGTG | tsx gene |
| T7_F | TAATACGACTCACTATAGGG | MCS pET28a(+) |
| T7-Terminator_R | GCTAGTTATTGCTCAGCGG | MCS pET28a(+) |
| pET28TEVSeqR | GAAATAAGGCTATGAGTCGC | MCS pET28a(+) |
| PBADseqF3 | ATCACGGCAGAAAAGTCCAC | MCS pBAD30 |
| PBADseqR3 | ACTCCCATCGGCGCTACGGC | MCS pBAD30 |
| ECygiVXF | AGCTGATCTCTAGATCGCAGGGAGGCAAAATGACAAACCTG | ygiV (*E. coli*) |
| ECygiVNF | AGCTGATCCATATGACAAACCTGACACTGGATG | ygiV (*E. coli*) |
| ECygiVH3R | AGCTGATCAGGGAAGCTTGAGTCAGGCATCACGCCAACGG | ygiV (*E. coli*) |
| ECygiWXF | AGCTGATCTCTAGAGGGAGTAATAAACATGAAAAAATTCGCAGCA | ygiW (*E. coli*) |
| ECygiWH3R | AGCTGATCAAGCTTCCCGGGAGCGGTAACAATTACGGATTTAC | ygiW (*E. coli*) |
| pUC/M13(-40) | GTTTTCCCAGTCACGAC | MCS pUC57 |

*GAAAACCTGTATTTTCAGGGC = TEV site
Abbreviations: DB, DNA binding domain; LBD, ligand-binding domain; MCS, multiple cloning site

**TableS11.** STM3175-constructs cloned in this work.

| Construct | Description | Primer combination |
|---|---|---|
| STM3175 | | 3175NdelF/3175BamHIR |
| STM3175-DB-$P_{114}$ | STM3175-N-terminal to $P_{114}$ | 3175NdelF/3175BamHIPR |
| STM3175-DB-$V_{133}$ | STM3175-N-terminal to Val133 | 3175NdelF/3175BamHIVR |
| STM3175-LBD-$P_{114}$ | STM3175-C-terminal from Pro114 | 3175NdeIPF/3175BamHIR |
| STM3175-LBD-$V_{133}$ | STM3175-C-terminal from Val133 | 3175NdeIVF/3175BamHIR |

**Abbreviations: DB, DNA binding domain; LBD, ligand-binding domain**
**STM3175 coding sequence:**
$M_0$NDLISAAYSERLRRVCDHIERHLDEPLSIEALSRMAHSSPFHFHRQFTTWSGLPLYRYIQWL-RLRRASWRLAFNPQDKVIDIALDAGFQNPESFTRAFKTAFGQSPRRFRQS$P_{114}$DWLAWHQRVPKLALQEQH$V_{133}$MDVKIVEFPPTRVAMLTHLGHPDKVNASAAKFIAWRRETGQSPIASS-QTFGIAWHDPQTTP-PAQFRFDICGSVRQPIAENDVGVVNSEIPGGRCAVVRHQGSLDSLPESVWYLFREWLPASGETPRDFPVFFQYLNFVHEVAEHELLTDIYLPL$R_{288}$

**Table S12:** Diffraction data collection, refinement, and validation statistics.

| **Dataset** | STM3175 |
|---|---|
| PDB entry | 7R3W |
| **Data Collection** | |
| Wavelength [Å] | 0.9184 |
| Temperature [K] | 100 |
| Space group | $C222_1$ |
| Unit Cell Parameters<br>a, b, c [Å]<br>α, β, γ [°] | <br>120.7, 244.9, 139.3<br>90.0, 90.0, 90.0 |
| Resolution [Å][a] | 30.00 – 3.60<br>(3.82 – 3.60) |
| Reflections [a]<br>Unique [a]<br>Completeness [%][a]<br>Multiplicity [a] | <br>24,093 (3,804)<br>99.5 (99.2)<br>5.6 (5.9) |
| Data quality [a]<br>Intensity [I/σ(I)] [a]<br>$R_{meas}$ [%][a,b]<br>$CC_{1/2}$ [a,c]<br>Wilson B value [$Å^2$] | <br>8.17 (0.98)<br>23.3 (196.9)<br>99.6 (49.6)<br>99.6 |
| **Refinement** | |
| Resolution [Å][a] | 30.00 - 3.60<br>(3.75 - 3.60) |
| Reflections [a]<br>Number<br>Test Set [%] | <br>23,959<br>5.0 |
| $R_{work}$ [%][a]<br>$R_{free}$ [%][a] | 26.8 (41.5)<br>31.5 (44.7) |
| Asymmetric Unit<br>Protein: Residues, Atoms | <br>285 (A), 2,330 (A), 282 (B), 2,306 (B)<br>282 (C), 2,306 (C), 287 (D), 2,346 (D) |
| Mean Temperature factors [$Å^2$][b]<br>All Atoms<br>Macromolecules | <br>134<br>127 (A), 131 (B)<br>147 (C), 130 (D) |
| RMSD from Target Geometry [d] | |

| | |
|---|---|
| Bond Lengths [Å] | 0.003 |
| Bond Angles [°] | 0.685 |
| **Validation Statistics** | |
| Ramachandran Plot [f] | |
| Residues in Allowed Regions [%] | 3.3 |
| Residues in Favored Regions [%] | 96.6 |
| Ramachandran plot *Z*-score [f] (RMSD) | |
| whole | -1.83 (0.22) |
| helix | -1.10 (0.19) |
| sheet | -0.68 (0.35) |
| loop | -1.40 (0.26) |
| MOLPROBITY Clashscore [g] | 10.1 |
| MOLPROBITY score [f] | 1.74 |

[a] data for the highest resolution shell in parenthesis
[b] $R_{meas}(I) = \sum_h [N/(N-1)]^{1/2} \sum_i \left| I_{ih} - <I_h> \right| / \sum_h \sum_i I_{ih}$, in which $<I_h>$ is the mean intensity of symmetry-equivalent reflections h, $I_{ih}$ is the intensity of a particular observation of h and N is the number of redundant observations of reflection h. (*26*)
[c] $CC_{1/2} = (<I^2> - <I>^2) / (<I^2> - <I>^2) + \sigma^2_\varepsilon$, in which $\sigma^2_\varepsilon$ is the mean error within a half-dataset. (*27*)
[d] RMSD – root mean square deviation
[e] calculated with PHENIX (*21*)
[f] calculated with MOLPROBITY (*28*)
[g] Clashscore is the number of serious steric overlaps (> 0.4 ) per 1,000 atoms (*28*)

**Table S13:** List of 2063 quantified proteins from Salmonella Typhimurium detected in wild type strain (n=5) and evolved T12 strain (n=4*). The mass spectrometry proteomics data have been deposited to the ProteomeXchange Consortium via the PRIDE partner repository with the dataset identifier PXD031944.
